# Protein manipulation using single copies of short peptide tags in cultured cells and in *Drosophila melanogaster*

**DOI:** 10.1101/2020.04.06.027599

**Authors:** M. Alessandra Vigano, Clara-Maria Ell, Manuela MM Kustermann, Gustavo Aguilar, Shinya Matsuda, Ning Zhao, Timothy J Stasevich, George Pyrowolakis, Markus Affolter

**Affiliations:** Growth and Development, Biozentrum, University of Basel, Klingelbergstrasse 70, CH-4056 Basel, Switzerland; Spemann Graduate School of Biology and Medicine (SGBM), Albert-Ludwigs University of Freiburg, 79104 Freiburg, Germany; Institute for Biology I, Faculty of Biology, ZBSA, Habsburgerstr. 49, Albert-Ludwigs University of Freiburg, 79104 Freiburg, Germany; CIBSS □ Centre for Integrative Biological Signalling Studies, Albert-Ludwigs University of Freiburg, 79104 Freiburg, Germany; Department of Biochemistry and Molecular Biology, Colorado State University, Fort Collins, CO 80523, USA

## Abstract

Cellular development and specialized cellular functions are regulated processes which rely on highly dynamic molecular interactions among proteins, distributed in all cell compartments. Analysis of these interactions and their mechanisms of action has been one of the main topics in cellular and developmental research over the last fifty years. Studying and understanding the functions of proteins of interest (POIs) has been mostly achieved by their alteration at the genetic level and the analysis of the phenotypic changes generated by these alterations. Although genetic and reverse genetic technologies contributed to the vast majority of information and knowledge we have gathered so far, targeting specific interactions of POIs in a time- and space-controlled manner or analyzing the role of POIs in dynamic cellular processes such as cell migration or cell division would require more direct approaches. The recent development of specific protein binders, which can be expressed and function intracellularly, together with several improvements in synthetic biology techniques, have contributed to the creation of a new toolbox for direct protein manipulations. We selected a number of short tag epitopes for which protein binders from different scaffolds have been developed and tested whether these tags can be bound by the corresponding protein binders in living cells when they are inserted in a single copy in a POI. We indeed find that in all cases, a single copy of a short tag allows protein binding and manipulation. Using *Drosophila*, we also find that single short tags can be recognized and allow degradation and relocalization of POIs *in vivo*.

## Introduction

A key question in cell and developmental biology is how the millions of protein molecules present in any given cell regulate cellular functions in a predictable and coordinated manner. Much of the work done in the past decades to study protein function in their *in vivo* setting has relied on the use of genetic and reverse genetic approaches which, combined with biochemical and structural studies, have been extremely successful in gaining insight into protein function (Housden et al., 2017; Wang et al., 2016). However, it turned out that any given protein can interact with many different partners, often in a location- or context-dependent fashion, in many cases regulated by specific posttranslational modifications. The complexity of protein-protein interactions has made it very difficult to decipher the manifold properties of any given protein of interest (POI) by using existing gain- and loss-of-function genetic studies. It would be desirable to have at hand a diversified toolbox to manipulate proteins directly in time and space in more controllable fashions.

Over the past few years, several novel approaches have opened up the way to specifically and directly manipulate the function of POIs in different ways in living cells or organisms and analyse the consequences of such manipulation on cellular or organismal level.

On the one hand, optogenetic tools have allowed users to manipulate proteins by fusing them to optically regulated modules using light as an inducer. These tools are mostly based on the properties of certain natural occurring photosensitive proteins to change their conformation or aggregation state in response to specific wavelengths (Tischer and Weiner, 2014). These proteins have been engineered into optogenetic systems to control neuronal activity (Rost et al., 2017), direct subcellular localization (Buckley et al., 2016; Niopek et al., 2016), turn protein functionality on or off (Bonger et al., 2014), promote gene expression or repression, or induce protein degradation and regulate cell signalling (Repina et al., 2017; Zhang and Cui, 2015).

Alternatively, chemically regulated modules can also be fused to POIs such that some of their functions (half-live, localization, etc.) can be manipulated (Banaszynski et al., 2006; Bonger et al., 2011; Chung et al., 2015; Czapinski et al., 2017; Natsume and Kanemaki, 2017; Natsume et al., 2016).

On the other hand, protein binders such as scFvs, nanobodies, DARPins, Affibodies, Monobodies and others have been used to directly target and manipulate POI in different cellular environments (extracellular or different intracellular compartments). These protein binders can be functionalized to allow the regulation of POIs in a desired manner. Using functionalized protein binders, POIs can be visualized, degraded, delocalized, or post-transcriptionally modified *in vivo* in order to learn more about the function of the POIs in cultured cells or in developing organisms (Aguilar et al., 2019a; Bieli et al., 2016; Harmansa and Affolter, 2018; Helma et al., 2015; Holliger and Hudson, 2005; Pluckthun, 2015; Prole and Taylor, 2019; Schumacher et al., 2018).

Several strategies allow to target and manipulate POIs *in vivo* via the use of protein binders. Binders against proteins can be isolated using existing platform and/or libraries, functionalized in a desired manner and expressed in cells or organisms upon transfection, viral transduction or from transgenes inserted into the genome (Dong et al., 2019; Dreier and Pluckthun, 2012; Fridy et al., 2014; McMahon et al., 2018; Moutel et al., 2016; Roder et al., 2017; Woods, 2019). Alternatively, binders against fluorescent tags can be used to manipulate a POI that has been fused to a fluorescent protein (FP). This strategy has the advantage that well validated FP binders are available, and that the fusion protein can be visualized during the process using confocal microscopy (Kaiser et al., 2014; Prole and Taylor, 2019). Ideally, and to minimize the potential perturbation of the POI, the latter could be tagged by a short peptide to which high affinity protein binders have been identified and characterized; this approach would allow the use of characterized and validated binders and results in minimal potential disturbance of the function of the POI. Multiple protein manipulation tools generated with nanobodies or DARPins directed towards FPs (Aguilar et al., 2019a; Beghein and Gettemans, 2017; Brauchle et al., 2014; Schumacher et al., 2018; Vigano et al., 2018) could be adapted in order to functionalize small tag binders.

Here, we have selected a number of existing short tag epitopes for which protein binders from different scaffolds have been reported in the last few years. We have tested whether these tags can be bound by the corresponding protein binders in living cells when they are inserted in a single copy in a POI. We indeed find that in most cases, a single copy of a short tag allows protein binding and manipulation. Using *Drosophila*, we show that single short tags can also be recognized *in vivo* in developing organisms and allow protein degradation and protein relocalization. Using combinations of these short tags and their corresponding, well-characterized binders will allow for many interesting protein manipulations with minimal functional interference and using validated reagents for POI binding and manipulation.

## Results

We wanted to investigate whether small tag binders (such as single chain fragment v (scFv) and nanobody (Nb)), which were shown to work *in vivo* as intrabodies, were able to bind a single short peptide tag inserted in proteins located in different cellular compartments.

We used transient transfection in HeLa cells as a model system to test the binding properties of these protein binders (Brauchle et al., 2014; Moutel et al., 2016; Vigano et al., 2018). We therefore generated mammalian expression constructs for the anti-GCN4 (SunTag) scFv (Tanenbaum et al., 2014), the anti-gp41(MoonTag) nanobody 2H10 (Boersma et al., 2019), the anti-HA scFvs frankenbodies (Zhao et al., 2019) and the anti-ALFA nanobody (Gotzke et al., 2019) fused to either sfGFP or mEGFP for intracellular visualization. All the binders were expressed under the control of the strong CMV promoter/enhancer (for a summary of binder constructs, see Figure 1). We next generated differently localized cellular “baits” containing one single copy of each tag fused to different proteins or protein domains for localization purposes, and to mCherry for visualization. Figure 1 shows a schematic representation of these baits and Suppl. Figure1a/Figure1b shows their subcellular localization upon transfection (see also Materials and Methods).

**Figure 1.**
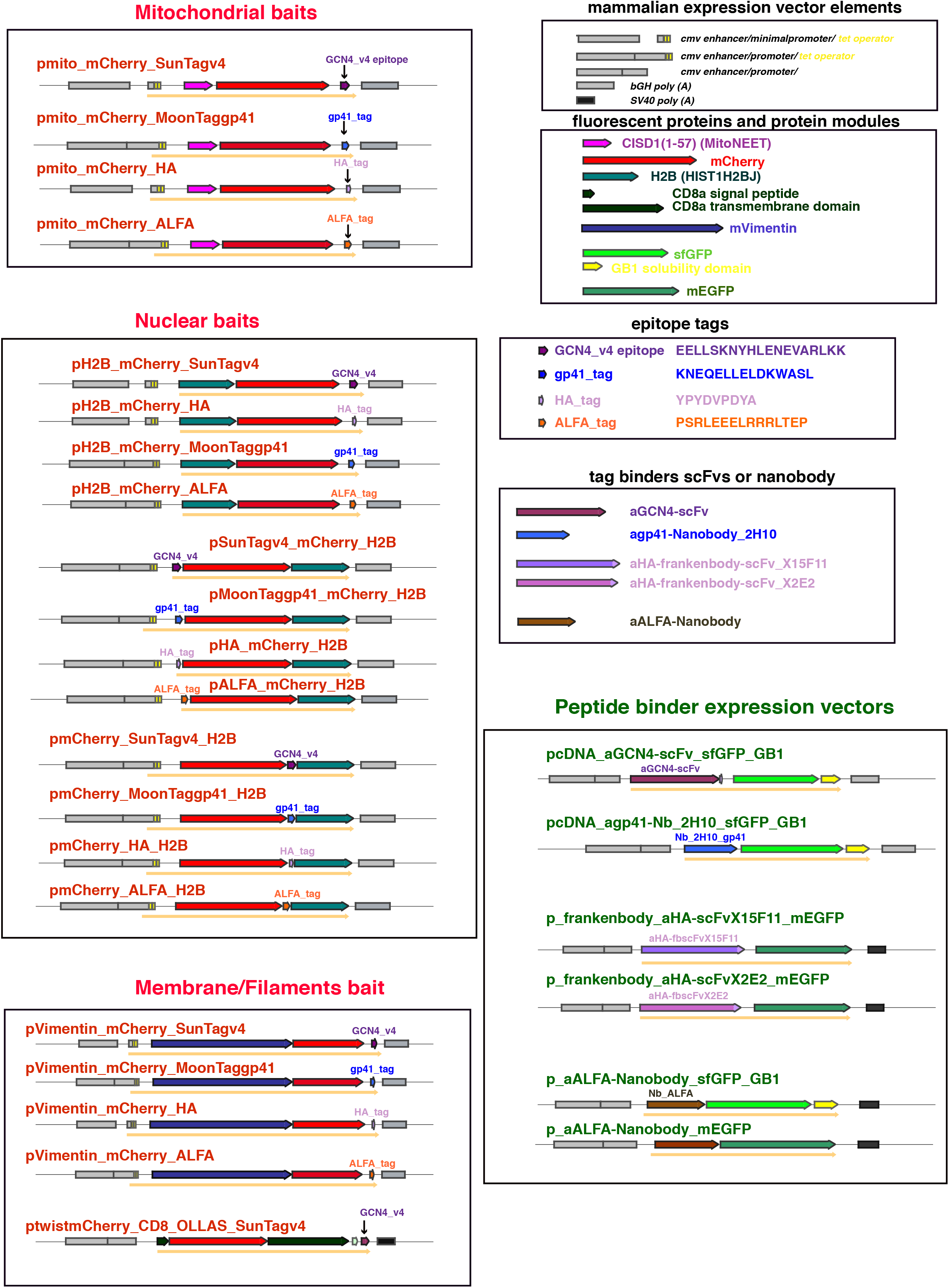
Schematic representation of the constructs. The transcriptional elements (enhancer, promoter and poly (A) adenylation) of the different mammalian expression vectors are depicted as grey filled boxes. The different protein coding modules are represented as coloured block arrows, while the resulting fusion protein is depicted as a solid orange arrow below the modules. Full maps and sequences are available upon request.

The mitochondrial baits were derived from the plasmid pcDNA4TO-mito-mCherry-10xGCN4_v4 (Tanenbaum et al., 2014), which encodes the N-terminal domain of the outer mitochondrial membrane protein MitoNEET (CISD1). This domain is N-terminally anchored to the outer membrane of the mitochondria and exposed to the cytoplasmic environment (Colca et al., 2004; Wang et al., 2017). This N-terminal domain of CISD1 is fused to mCherry and to either one of the tags we tested (GCN4- v4: 19aa; gp41: 15aa; HA: 9aa; ALFA: 15aa) in a single copy in the C-terminal position. The expression pattern of these different mitochondrial constructs in transfected cells were very similar, and most of the mitochondria around the nuclei were decorated by the mCherry protein, with almost no expression visible in the cytoplasm but possible expression in other internal membrane compartments. We also noted a slightly different pattern of expression for the pmito_mCherry_MoonTaggp41 (Suppl. Figure1a, panels of B); in this case, the mitochondria appeared less rounded and more filamentous; furthermore, the cytoplasmic mCherry signal was slightly stronger. A stronger cytoplasmic signal was also observed for pmito_mCherry_ALFA (Suppl. Figure 1a, panels of D).

The nuclear baits were based on histone H2B (H2BC11) fused to mCherry either at the N- (pH2B_mCherry) or C-terminus (pmCherry_H2B) and with the individual tags located at the N-terminus (pTag_mCherry_H2B), between mCherry and H2B (pmCherry_Tag_H2b) or at the C-terminus (pH2B_mCherry_Tag). As seen in Suppl. Figure1b, all these nuclear baits were located exclusively to the nucleus upon transient expression, although some appeared more concentrated in nucleoli or unspecific nuclear bodies, irrespective of the position of the H2B or the nature or position of the peptide tag (see pH2B_mCherry_SunTagv4, panels of A; pH2B_mCherry_HA, panels of C; pmCherry_ALFA_H2B, panels of H; pSunTagv4_mCherry_H2B, panels of I; pHA_mCherry_H2B, panels of K; pALFA_mCherry_H2B, panels of L; pmCherry_H2B, panels of M). The different localizations in the nucleus might be due to an accumulation in particular sub-nuclear structures for coping with the overexpression (Amer-Sarsour and Ashkenazi, 2019; Rekulapally and Suresh, 2019) or might reflect the different localization of the H2B fusion protein during the cell cycle phases (Duronio and Marzluff, 2017; Kurat et al., 2014; Romeo and Schumperli, 2016). Moreover, it could also reflect the rapid turnover of the histone H2B specifically in chromatin domains with high transcriptional activity (Kimura and Cook, 2001).

We also generated a bait with the leader sequence and the transmembrane domain of the mouse CD8 protein fused to mCherry and containing both the OLLAS (Park et al., 2008) and the GCN4-v4 tags. In *Drosophila melanogaste*r, this construct arrangement was shown to be inserted into the plasma membrane, exposing the mCherry moiety in the extracellular space and the domains at the C-terminus of CD8 (in this case the two small tags) at the cytoplasmic side of the membrane (Harmansa et al., 2015). In the mammalian system, fusion constructs to the CD8 protein domains have been used, for example to study trans Golgi vesicular transport (Nickel et al., 1998; Pascale et al., 1992a; Pascale et al., 1992b). As shown in Suppl. Figure1a, panels of I, expression of the pTwist_mCherry_CD8_OLLAS_SunTagv4 construct in HeLa cells, painted not only the plasma membrane, but other membranous and filamentous structures inside the cytoplasm with mCherry.

The last subcellular bait was a fusion between the mouse Vimentin protein, mCherry and one copy of each peptide tag at the C-terminus (Gotzke et al., 2019). As shown in Suppl. Figure1a, these constructs reflected the expression of Vimentin in the intermediate filaments of the transfected cells, although the filaments painted by the construct containing the HA tag appeared slightly different, shorter, thicker and with a sort of punctuate structure (panels of G).

### SunTag

The SunTag system was developed by (Tanenbaum et al., 2014) to visualize protein expression and translation in high resolution fluorescence imaging. The tag (v1) is an epitope derived from the yeast amino acid starvation-responsive transcription factor GCN4, subsequently optimized (v4) for binding to a previously characterized scFv with specific intracellular expression (Worn et al., 2000). As shown in Figure 2 (panels of A) or Suppl. Figure2 (panels of A), anti-GCN4scFv was uniformly distributed both in the cytoplasm and in the nucleus of the transfected cells, with a stronger green signal (from the sfGFP fusion partner) in the nucleus. This nuclear signal was not entirely overlapping with the Hoechst staining (which highlights mostly the DNA), indicating free diffusion of the scFv in the nucleoplasm. Occasionally, we observed some aggregation/accumulation in some unidentified granular structures in the cytoplasm, possibly due to the high level of expression of the construct.

**Figure 2.**
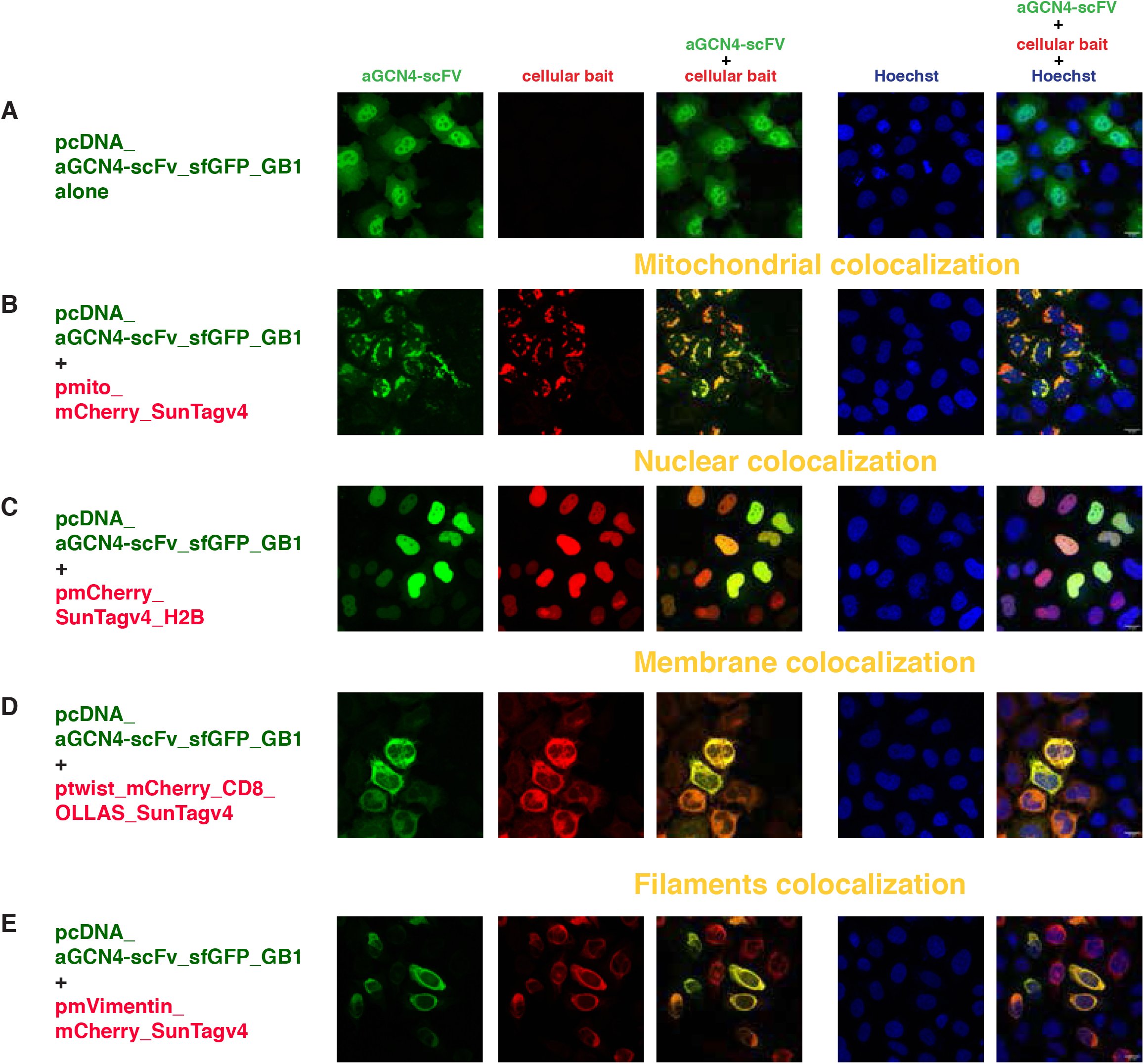
Intracellular binding of anti-GCN4 scFv (SunTag system) Confocal images of HeLa cells transiently transfected with **(A)** pcDNA_aGCN4- scFv_sfGFP_GB1 alone; the combination of pcDNA_aGCN4-scFv_sfGFP_GB1 and **(B)** pmito_mCherry_SunTagv4,**; (C)** pmCherry_SunTagV4_H2B; **(D)** ptwist_mCherryCD8_OLLAS_SunTagV4; **(E)** pmVimentin_mCherry_SunTagV4. The first column represents the GFP channel (green), the second column is the mCherry channel (red), the third column is the overlay of the two channels, showing the colocalization (indicated in yellow) of the antiGCN4 scFvs with the respective mitochondrial **(B)**, nuclear **(C)**, membrane **(D**) and filaments **(E)** baits; the fourth column represents the nuclear Hoechst staining (blue) and the fifth column is the merge of all three channels (with the scale bar in white (15 μm) on the bottom right corner). Images were taken 24 hours post transfection. Transfected constructs are indicated at the left of each row and the single and merge channels are indicated at the top of the respective columns. The figures are from a representative experiment, performed at least three times.

Coexpression of anti-GCN4scFv with the mitochondrial bait carrying a single copy of the SunTag epitope v4 changed significantly the distribution of the anti-GCN4scFv, mostly relocalizing it to the outer mitochondrial membrane (Figure. 2, panels of B). The pmito_mCherry_SunTagv4 did not change its localization at the mitochondria of the transfected cells when expressed with anti-GCN4scFv (compare expression either alone (Suppl. Figure1a, panels of A) or in combination with anti-GCN4scFv (Figure 2, panels of B).

It has to be noted that not all the anti-GCN4scFv molecules were recruited to the mitochondria, as seen by residual sfGFP signal in the cytoplasm, presumably because of the limited number of CISD1 binding partners at the mitochondrial surface. Varying the ratio of the transfected DNAs did not change the amount of anti-GCN4scFv observed at the mitochondria (data not shown).

Importantly, we did not observe any colocalization of anti-GCN4scFv with similar mitochondrial baits carrying one copy of either the unrelated HA epitope tag (Suppl. Figure 3, panels of A) or the gp41 epitope (MoonTag) (Suppl. Figure 3, panels of B), suggesting that one copy of the SunTag located at the C-terminus of the mito-mCherry fusion construct was indeed sufficient to specifically bind and recruit the majority of the anti-GCN4scFv to the outer mitochondrial membrane.

We also generated a mitochondrial bait containing one copy of the original GCN4 peptide tag v1 (Tanenbaum et al., 2014) and observed the same recruitment to the outer mitochondrial membrane of the anti-GCN4scFv (data not shown).

We next tested for nuclear colocalization with three different nuclear baits, all based on the histone protein H2B but with a single copy of the SunTag epitope in different positions (see Figure 1). Cotransfection of these nuclear baits with the anti-GCN4scFv (Figure 2, panels of C, Suppl. Figure 2, panels of B and C) clearly showed nuclear accumulation of the scFv with a nearly complete overlap of the mCherry and GFP signal in the nuclei of transfected cells, including the nucleoli/nuclear bodies. The residual GFP signal in the cytoplasm remained barely detectable. Also in these nuclear relocalization experiments, one single copy of the SunTag v4 epitope, in all the positions tested, was sufficient to bind and recruit the scFv to the nuclear compartments.

In the case of cotransfection of the anti-GCN4scFv with nuclear baits carrying the HA tag (Suppl. Figure 3, panels of C, D and E), the MoonTag (Suppl. Figure 3, panels of F), the ALFA tag (Suppl. Figure 3, panels of G) or no tag (pmCherryH2B) (Suppl. Figure 3, panels of H), we observed a partial overlap of the GFP and mCherry signal, especially in the nuclear bodies, but the majority of the anti-GCN4scFv was still visible in the cytoplasm and in the nucleoplasm, with a cellular localization very similar to the one observed in the absence of any bait. The strongest “cross reactivity” was observed with ALFA tag, maybe due to a certain similarity of the two tags (the conserved EEL stretch might be sufficient for low affinity binding). Then, we tested the binding and localization of the anti-GCN4scFv in the presence of the CD8 “membrane” construct. As mentioned above, ptwist _mCherry_CD8_OLLAS_SunTagv4 localized both at the plasma membrane and at other filamentous structures associated with internal membranes of the transfected cells (shown in Suppl. Figure 1a, panels I). Its cellular localization was not changed when cotransfected with the anti-GCN4scFv, but it was able to bind and recruit the scFv, as illustrated by the almost complete overlap of the GFP and mCherry signal (Figure 2, panels of D.)

Finally, when tested in cotransfection with the Vimentin_mCherry_SunTagv4 bait, we observed an almost complete relocalization of the anti-GCN4scFv to the intermediate filaments revealed by the mCherry signal (Figure 2, panels of E), supporting an efficient binding *in vivo* of the anti-GCN4scFv to yet another subcellular compartment exposing a single copy of the SunTag. We also confirmed no binding of the anti-GCN4scFv to a coexpressed Vimentin bait with the MoonTaggp41 (Suppl. Figure 3, panels of I) and some cross reactivity with the Vimentin bait containing the ALFA tag (Suppl. Figure 3, panels of J).

### MoonTag

The recent development of the MoonTag system (Boersma et al., 2019) prompted us to test this system in a similar way to the SunTag. MoonTag is based on the epitope from the membrane-proximal external region of the human HIV-1 envelope glycoprotein subunit gp41 and its nanobody binder 2H10. We therefore generated the mitochondrial, the Vimentin and the nuclear baits with a single copy of the MoonTag epitope and cloned the Nanobody anti-gp41 2H10 (anti-gp41Nb), fused to sfGFP_GB1, in a CMV promoter/enhancer expression vector (Figure 1).

The nuclear colocalization assay confirmed a very efficient binding and recruitment of the anti-gp41Nb to the nuclei with only one copy of the MoonTag in all the positions tested (Figure 3, panels of C, Suppl. Figure 2, panels of E and F); furthermore, no GFP signal was detected in the cytoplasm. Expression of the anti-gp41Nb alone resulted in rather uniform distribution of the protein in the cytoplasm (Figure 3, panels of A; Suppl. Figure 2, panels of D) and, as observed for the anti-GCN4scFv, with a stronger signal in the nuclei. We never observed any aggregation, possibly reflecting the better solubility of the nanobody than the scFv and confirming its good intracellular expression.

**Figure 3.**
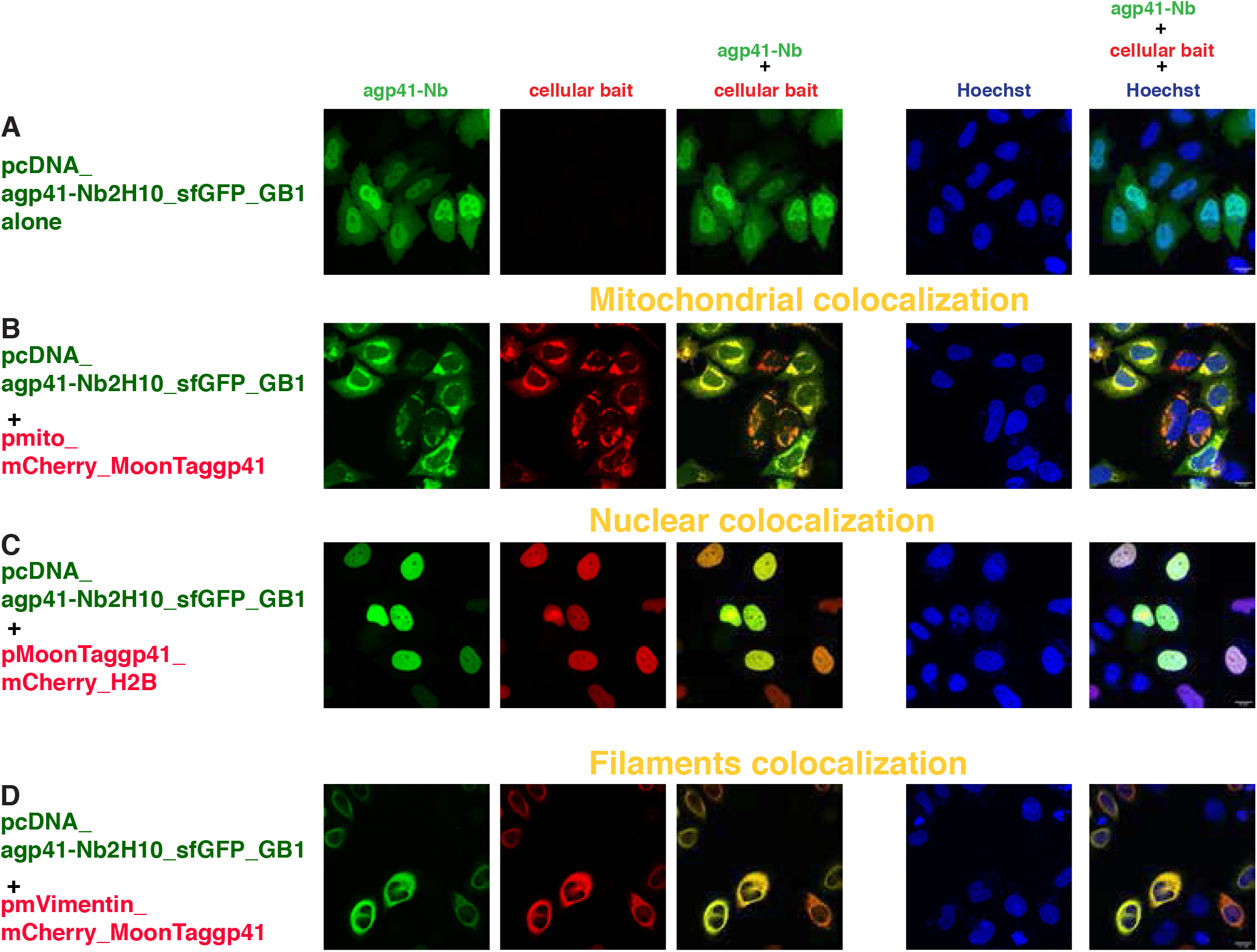
Intracellular binding of anti-gp41 Nanobody (MoonTag system) Confocal images of HeLa cells transiently transfected with **(A)** pcDNA_agp41- Nb2H10_sfGFP_GB1 alone; the combination of pcDNA_ agp41- Nb2H10_sfGFP_GB1 and **(B)** pmito_mCherry_MoonTaggp41; **(C)** pMoonTaggp41_mCherry_H2B; **(D)** pmVimentin_mCherry_MoonTaggp41. The first column represents the GFP channel (green), the second column is the mCherry channel (red), the third column is the overlay of the two channels, showing the colocalization (indicated in yellow) of the antigp41 nanobody with the respective mitochondrial **(B)**, nuclear **(C**) and filaments **(D**) baits; the fourth column represents the nuclear Hoechst staining (blue) and the fifth column is the merge of all three channels (with the scale bar in white (15 μm) on the bottom right corner). Images were taken 24 hours post transfection. Transfected constructs are indicated at the left of each row and the single and merge channels are indicated at the top of the respective columns. The figures are from a representative experiment, performed at least three times.

When the anti-gp41Nb was cotransfected with the nuclear bait carrying no tag (pmCherry-H2B) (Suppl. Figure 4, panels of F) or pH2B_mCherry_SunTagv4 (Suppl. Figure 4, panels of E), we also observed some overlapping GFP signal in the nuclear bodies with strong accumulation of the mCherry signal, but the majority of the GFP signal was uniformly distributed within the nucleus and the cytoplasm, where no mCherry signal was detected. As observed with the anti-GCN4scFv in the similar combination set up, these results are indicative of no binding or active recruitment by the nuclear baits with a different tag or with no tag.

In cotransfection experiments with the anti-gp41Nb and the mitochondrial bait carrying one copy of the MoonTag (Figure 3, panels of B), we observed both binding and recruitment of the anti-gp41Nb to the outer mitochondrial membrane of the transfected cells, although there was some detectable GFP signal in the cytoplasm and the nucleus.

Cotransfection of the anti-gp41Nb with mitochondrial baits either containing the HA (Suppl. Figure 4, panels of A), the SunTagv4 (Suppl. Figure 4, panels of B) or the ALFA tag (Suppl. Figure 4, panels of C) also showed a very partial overlap of the GFP and the mCherry signals, mostly with HA; however, most of the GFP signal remained uniformly distributed in the cytoplasm and the nucleus (especially with the pmito_mCherry_SunTagv4), with no indication of binding or active recruitment.

The colocalization of the anti-gp41Nb to the intermediate filaments was also very prominent (Figure 3, panels of D), indicating a very efficient binding and recruitment to these structures by Vimentin carrying one copy of the MoonTag. Furthermore, we did not observe any cross reactivity with Vimentin carrying the SunTagv4 (Suppl. Figure 4, panels of G) or the ALFA tag (Suppl. Figure 4, panels of H).

### HA tag

The HA peptide derived from the influenza virus hemagglutinin has been extensively used in biochemical studies due to the availability of high-affinity monoclonal antibodies (Field et al., 1988; Wilson et al., 1984). Recently, two different anti-HAscFvs derived from the monoclonal anti-HA antibody 12CA5 were generated and called frankenbodies (Zhao et al., 2019). The two frankenbodies anti-HA- scFvX15F11 and anti-HA-scFvX2E2 were made by grafting the complementarity determining regions (CDRs) of the 12CA5 monoclonal antibody into two different scFv scaffolds with a demonstrated solubility *in vivo*. We tested the function of these two anti-HAscFvs as intrabodies for their binding to a single copy of the HA epitope embedded in the same cellular baits as developed analogously for the SunTag and MoonTag systems (see Figure1).

The expression pattern of the two frankenbodies in the single transfection conditions in the absence of any bait was somewhat similar and equivalent to the intracellular distribution of the anti-GCN4scFv; uniform expression in both nucleus and cytoplasm, with stronger GFP signal in the nucleoplasm (see Figure 4, panels of A and Suppl. Figure 5, panels of A for p_frankenbody_anti-HA scFvX15F11_mEGFP and p_frankenbody_anti-HA scFvX2E2_mEGFP, respectively), confirming the expression of these scFvs reported by Zhao et al in a different cell line (U2OS)(Zhao et al., 2019).

**Figure 4.**
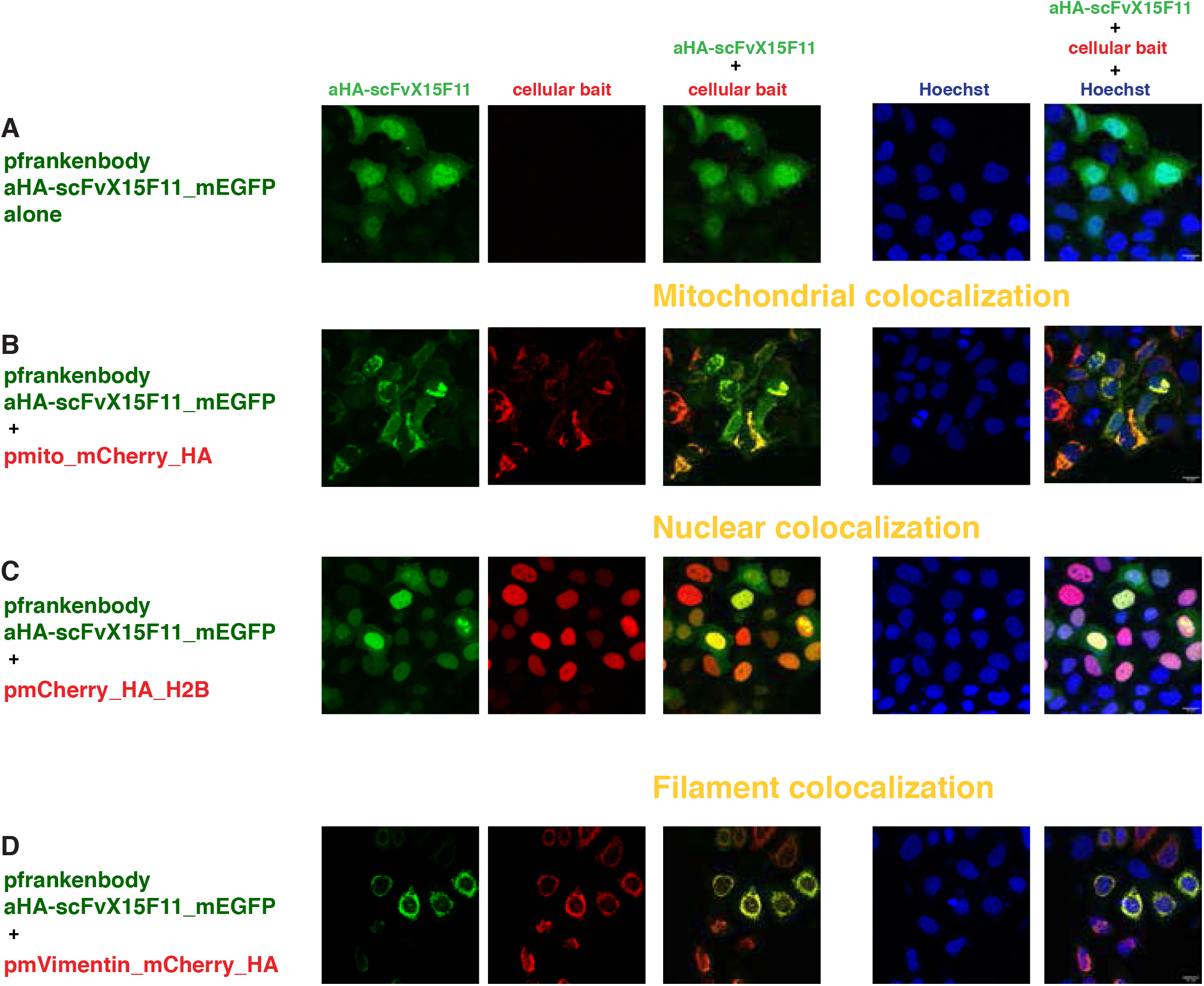
Intracellular binding of anti-HAscFv Frankenbody X15F11 (HA system) Confocal images of HeLa cells transiently transfected with **(A)** p_frankenbody_aHA- scFvX15F11_mEGFP alone; the combination of p_frankenbody_aHA- scFvX15F11_mEGFP and **(B)** pmito_mCherry_HA; **(C)** pmCherry_HA_H2B; **(D)** pmVimentin_mCherry_HA. The first column represents the GFP channel (green), the second column is the mCherry channel (red), the third column is the overlay of the two channels, showing the colocalization (indicated in yellow) of the antiHAscFv frankenbody with the respective mitochondrial **(B)**, nuclear **(C)** and filaments **(D**) baits; the fourth column represents the nuclear Hoechst staining (blue) and the fifth column is the merge of all three channels (with the scale bar in white (15 μm) on the bottom right corner). Images were taken 24 hours post transfection. Transfected constructs are indicated at the left of each row and the single and merge channels are indicated at the top of the respective columns. The figures are from a representative experiment, performed at least three times.

Cotransfection with the mitochondrial bait containing one copy of the HA epitope (p_mito_mCherry_HA) showed significant recruitment to the outer mitochondrial membrane of both frankenbodies (Figure 4, panels of B; Suppl. Figure 5, panels of B;). Although this kind of assay is not quantitative, we have the impression that the fraction of scFvs, which is detected in the cytoplasm or nucleoplasm (the GFP signal), is higher for anti-HA scFvX15F11 than for anti-HA scFvX2E2. The residual GFP signal not localizing at the mitochondrial membrane was also higher for these anti-HAscFvs than the anti-GCN4scFv signal in equivalent cotransfection conditions (Figure 2, panels of B). This may reflect a lower binding affinity of the scFvs to the HA epitope and consequently a lower efficiency of recruitment with only one epitope copy; this interpretation is in agreement with the lower signal to noise ratio of the Mito_mCherry_1xHA versus Mito_mCherry_smHA, containing 10xHA, reported by Zhao (Zhao et al., 2019).

We also tested the frankenbodies in mitochondrial colocalization experiments with mitochondrial baits containing the other tags, pmito_mCherry SunTagv4 (Suppl. Figure 6, panels of A and B) and pmito_mCherry_MoonTaggp41 (Suppl. Figure 6, panels of C and D). We did observe some partial colocalization with all the mitochondrial constructs and scFvs, with overlapping GFP and mCherry signals of different intensity and patterns in each combination of plasmids.

We next tested whether the nuclear baits containing one copy of the HA epitope positioned in different parts of the fusion proteins pHA_mCherry_H2B, pmCherry_HA_H2B and pH2B_mCherry_HA (Figure 1) were sufficient to actively bind and recruit the frankenbodies to chromatin. As shown in Figure 4, panels of C and Suppl.Figure 5, panels of A’and B’ for p_frankenbody_aHA scFvX15F11_mEGFP (and Suppl. Figure 5, panels of C,D and E for p_frankenbody_aHA scFvX2E2_mEGFP, there was a clear nuclear colocalization under all the condition tested, with a higher efficiency for the anti-HAscFvX2E2 than the anti-HAscFvX15F11, as judged from the residual GFP signal in the cytoplasm.

When we cotransfected the anti-HAscFvs with the pmCherry_H2B nuclear bait with no tag, we observed some overlapping GFP and mCherry signal in the nucleoli/nuclear bodies (as seen with others protein binders anti-GCN4 and anti-gp41), but the majority of the GFP signal was in the cytoplasm/nucleoplasm of transfected cells, as shown in Suppl. Figure 7, panels of A and B. In cotransfection experiments with nuclear bait containing the ALFA tag (Suppl. Figure 7, panels of C and D), we also observed minimal overlap of the mCherry and mEGFP signals in the nuclei. Hence, in the case of nuclear colocalization, one copy of the HA epitope, regardless of the insertion position, appeared sufficient to bind and recruit the frankenbodies to the nucleus, although somewhat less efficiently than anti-gp41Nb or anti-GCN4scFv counterparts (as judged from the residual GFP signal in the cytoplasm).

We also did an extra control with the p_frankenbody_anti-HA scFvX15F11_mEGFP to exclude any unspecific interaction with the unrelated bait ptwist _mCherry_CD8_OLLAS_SunTagv4. As shown in Suppl. Figure 6 panels E, we did not see any colocalization of the two constructs.

Cotransfection experiments of the two frankenbodies with Vimentin_mCherry_HA (Figure 4, panels of D, Suppl. Figure 5, panels of F) confirmed the binding to a single copy of the epitope in cultured cells, although we observed a higher residual GFP signal both in the cytoplasm and the nucleoplasm. Furthermore, as mentioned earlier, the expression of the pVimentin_mCherry_HA, either alone (Suppl. Figure 1a, panels of H) or with the anti-HA-scFvs, was significantly different from the intermediate filaments painted by the Vimentin_mCherry-SunTag or MoonTag, indicating a possible “disruption” of the filament structure of this particular Vimentin fusion protein. Nevertheless, the two anti-HAscFvs were able to bind to this HA bait specifically and they did not show binding with Vimentin_mCherry_ALFA (Suppl. Figure 7, panels of E and F).

### ALFA tag

Recently, Götzke et al developed a system they called ALFA tag, which is based on a short synthetic linear tag and its nanobody binder (Gotzke et al., 2019). We decided to test this new system as well, since we reasoned that nanobodies might be more versatile than scFvs as protein binders *in vivo* (see Discussion). Therefore, we generated the expression vector for the anti-ALFA nanobody fused either to sfGFP_GB1 or mEGFP and the same mitochondrial and nuclear baits carrying one copy of the ALFA tag in different positions (Figure 1).

We confirmed that the anti-ALFA nanobody is well expressed in transfected cells (Figure 5, panels of A, Suppl. Figure 8, panels of A and Suppl. Figure 9a panels of A and B), as reported by Götzke (Gotzke et al., 2019). In the case of anti-ALFA nanobody fused to mEGFP, we occasionally observed some minor aggregation (data not shown), but the expression pattern of both anti-ALFA nanobodies fused to sfGFP or mEGFP was very similar to the other binders tested above.

**Figure 5.**
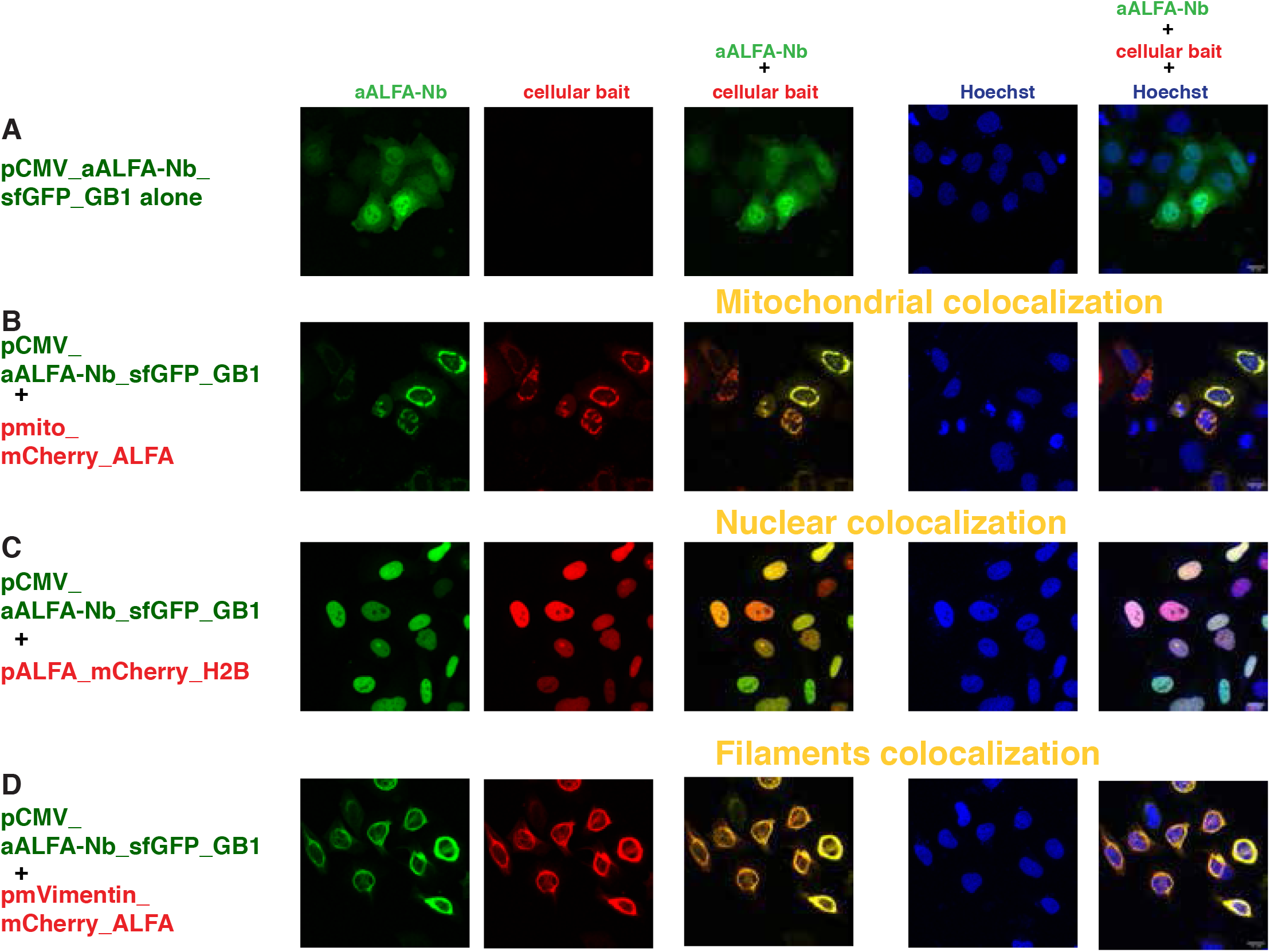
Intracellular binding of anti-ALFA Nanobody (ALFA tag system) Confocal images of HeLa cells transiently transfected with **(A)** pCMV_aALFA- Nb_sfGFP_GB1 alone; the combination of pCMV_aALFA-Nb _sfGFP_GB1 and **(B)** pmito_mCherry_ALFA; **(C)** pALFA_mCherry_H2B; **(D)** pmVimentin_mCherry_ALFA. The first column represents the GFP channel (green), the second column is the mCherry channel (red), the third column is the overlay of the two channels, showing the colocalization (indicated in yellow) of the anti-ALFA nanobody with the respective mitochondrial **(B**), nuclear (**C**) and filaments (**D**) baits; the fourth column represents the nuclear Hoechst staining (blue) and the fifth column is the merge of all three channels (with the scale bar in white (15 μm) on the bottom right corner). Images were taken 24 hours post transfection. Transfected constructs are indicated at the left of each row and the single and merge channels are indicated at the top of the respective columns. The figures are from a representative experiment, performed at least three times.

In cotransfection experiments with the pmito_mCherry_ALFA bait, the binding and recruitment to the outer mitochondrial membrane of the anti-ALFA nanobody was very efficient (Figure 5, panels of B, Suppl. Figure 8, panels of B), and the residual cytoplasmic signal was virtually negligible. Control experiments with mitochondrial baits containing the MoonTaggp41 (Suppl. Figure 9b, panels of A and B) revealed no cross reactivity.

The nuclear colocalization was also very efficient with all the nuclear baits tested, irrespective of the position of the ALFA tag (Figure 5, panels of C, Suppl. Figure 8, panels of A’ and B’ for a-ALFAnanobody_sfGFP_GB1 and Suppl. Figure 8, panels of C, D and E for a-ALFAnanobody-mEGFP). Control experiments with nuclear baits containing different tags (Suppl. Figure 9a, panels of C,D,E and F) or no tag (Suppl. Figure 9a, panels of G and H) showed a detectable nuclear colocalization with the pH2B_mCherry_SunTagv4 and a partial overlap with the pmCherry_H2B signal, although most of the nanobodies’ signals was still detectable in the cytoplasm.

Finally, we tested binding and recruitment to the intermediate filaments with a Vimentin_mCherry bait carrying one copy of the ALFA tag at the C-terminus. As reported with a similar Vimentin construct, but with the ALFA tag at the N-terminus (and without fluorescent protein) (Gotzke et al., 2019), we observed excellent colocalization of the mCherry and GFP signals (Figure 5, panels of D, Suppl. Figure 8, panels of F). Furthermore, we did not observe any cross reactivity with Vimentin-SunTagv4 (Suppl. Figure 9b, panels of C and D) or MoonTaggp41 (Suppl. Figure 9b, panels of E and F).

### Binding and manipulation of single HA tagged proteins *in vivo*

We next addressed whether single tagged POIs can be recognized and manipulated by the respective binders *in vivo*. We used *Drosophila* as a test system and focused on the HA tag, as this epitope is widely used to mark proteins in the *Drosophila* research field. We generated transgenic flies expressing an aHA-scFvX15F11_mEGFP fusion protein under the control of the UAS/GAL4 system. When expression was activated in salivary glands, the GFP signal was ubiquitously distributed throughout the cell. Similarly to the cotransfection experiments, GFP levels were slightly increased in the nuclei (Figure 6, panels of A). Co-expression of nuclear-localized Histone H2Av carrying a single HA tag at the C-terminus (H2Av-Flag-1xHA) resulted in a strong accumulation of the aHA-scFvX15F11_mEGFP in the nucleus (Figure 6, panels of B and C). Similar to the corresponding cell culture experiment (Figure 4 and Suppl. Figure 5), the cytoplasmic pool of aHA-scFvX15F11_mEGFP was reduced but not completely depleted. To address whether the efficacy of nuclear translocation of aHA-scFvX15F11_mEGFP might depend on the number of epitope tag copies in the POI, we used *Drosophila* Histone H4 carrying three HA copies at its C-terminus as nuclear bait (H4-3xHA). Indeed, using the same experimental setting, co-expression of aHA-scFvX15F11_mEGFP with H4-3xHA (Suppl. Figure 10) resulted in strong accumulation of the eGFP signal in the nucleus and its complete depletion from the cytosol. Cumulatively, these findings suggest that HA-binders can be utilized for efficient binding of proteins *in vivo*, with the efficiency being a function of the copy-number of HA epitopes in the POI.

**Figure 6.**
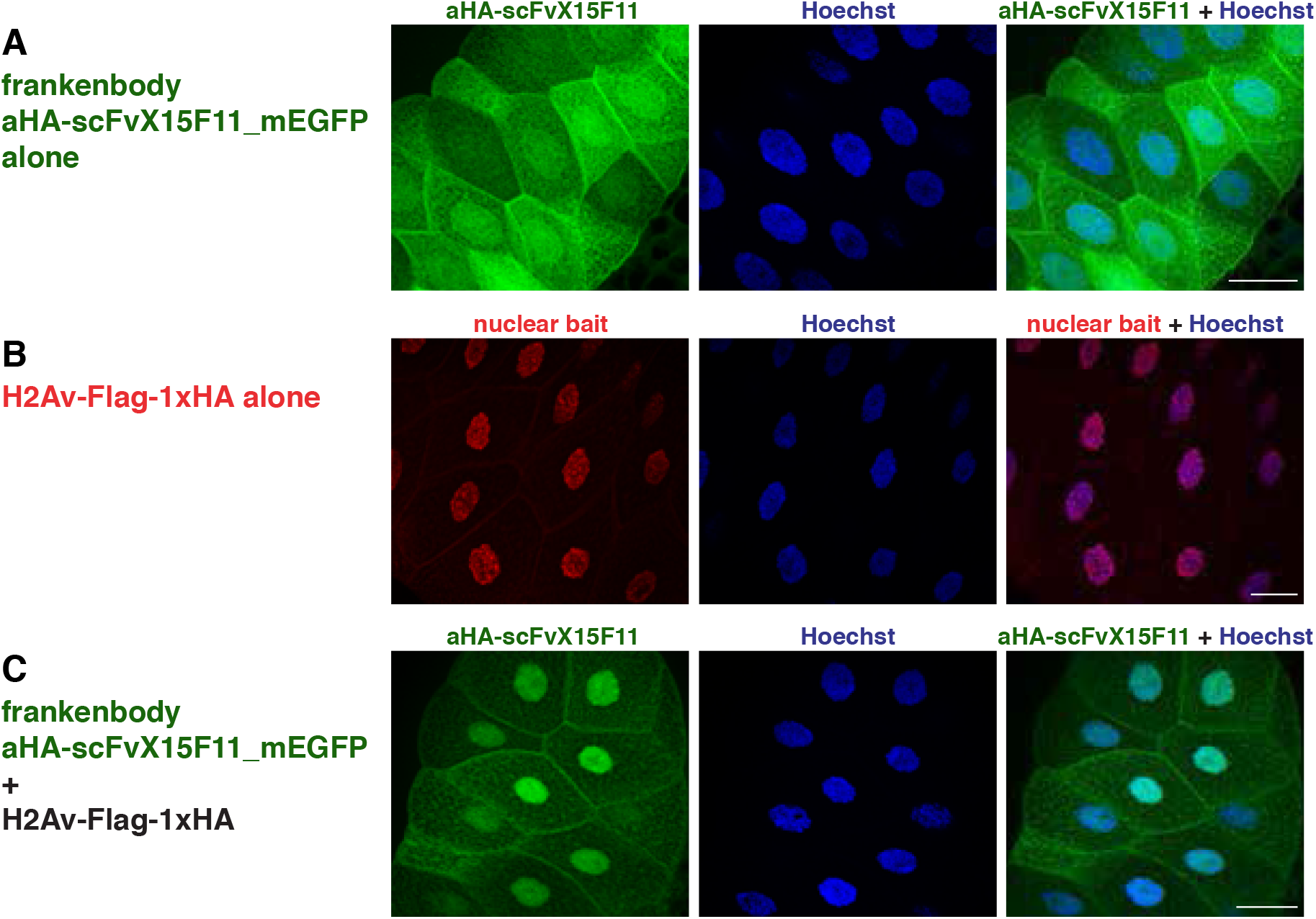
Intracellular binding of anti-HAscFv Frankenbody X15F11 (HA system) *in vivo*. Confocal images of *Drosophila* larval salivary glands expressing frankenbody_aHA- scFvX15F11_mEGFP alone (**A**), the nuclear bait H2Av-Flag-1xHA alone (**B**), or a combination of frankenbody_aHA-scFvX15F11_mEGFP and H2Av-Flag-1xHA (**C**). The first column represents the GFP channel (green, **A** and **C**) or the anti-Flag staining channel (red, **B**). The second column represents the nuclear Hoechst staining (blue) and the third column is the merge of the two respective channels. Scale bars are 50 µm. Salivary glands were obtained from third instar *Drosophila* larvae expressing the constructs indicated at the left of each row. Single and merge channels are indicated at the top of the respective channel.

We also tested whether single-tagged POIs can be inactivated by functionalized protein binders. Previous work established a tool, deGradFP, allowing for ubiquitin/proteasome degradation of GFP-tagged proteins using a nanobody against GFP (Caussinus et al., 2011). In this system, a single-domain antibody fragment against GFP (vhhGFP4) was used to replace the substrate specificity domain of the Drosophila E3 component Slmb generating an E3 ligase complex that is directed against GFP and GFP-tagged proteins. We modified the deGradFP tool by replacing the vhhGFP4 domain with aHA-scFvX15F11 to generate deGradHA and tested its activity towards HA-tagged proteins in transgenic flies. As a POI we used Yorki (Yki, *Drosophila* YAP/TAZ), a transcriptional co-activator that is regulated through phosphorylation by the Hippo signalling pathway to control cell proliferation and organ size (Huang et al., 2005). In the construct used, YkiS168A-HA-eGFP, Yki contains a point mutation that renders the protein hyperactive in promoting organ growth (Oh and Irvine, 2008). In addition, the protein contains a C-terminal single HA tag followed by GFP. As shown before, transgenic flies expressing YkiS168A- HA-eGFP using an eye-specific driver display massive tissue overgrowth (Figure 7, panels A and B)(Oh and Irvine, 2008). This phenotype can be completely reversed by co-expressing deGradHA (Figure 7, panel C) or deGradFP (not shown) but not by co-expression of an unrelated protein (ß-galactosidase (lacZ); Figure 7, panel D), the later excluding titration effects of the UAS/GAL4 system. Thus, the deGradHA tool can efficiently inactivate proteins carrying single HA epitope tag.

**Figure 7.**
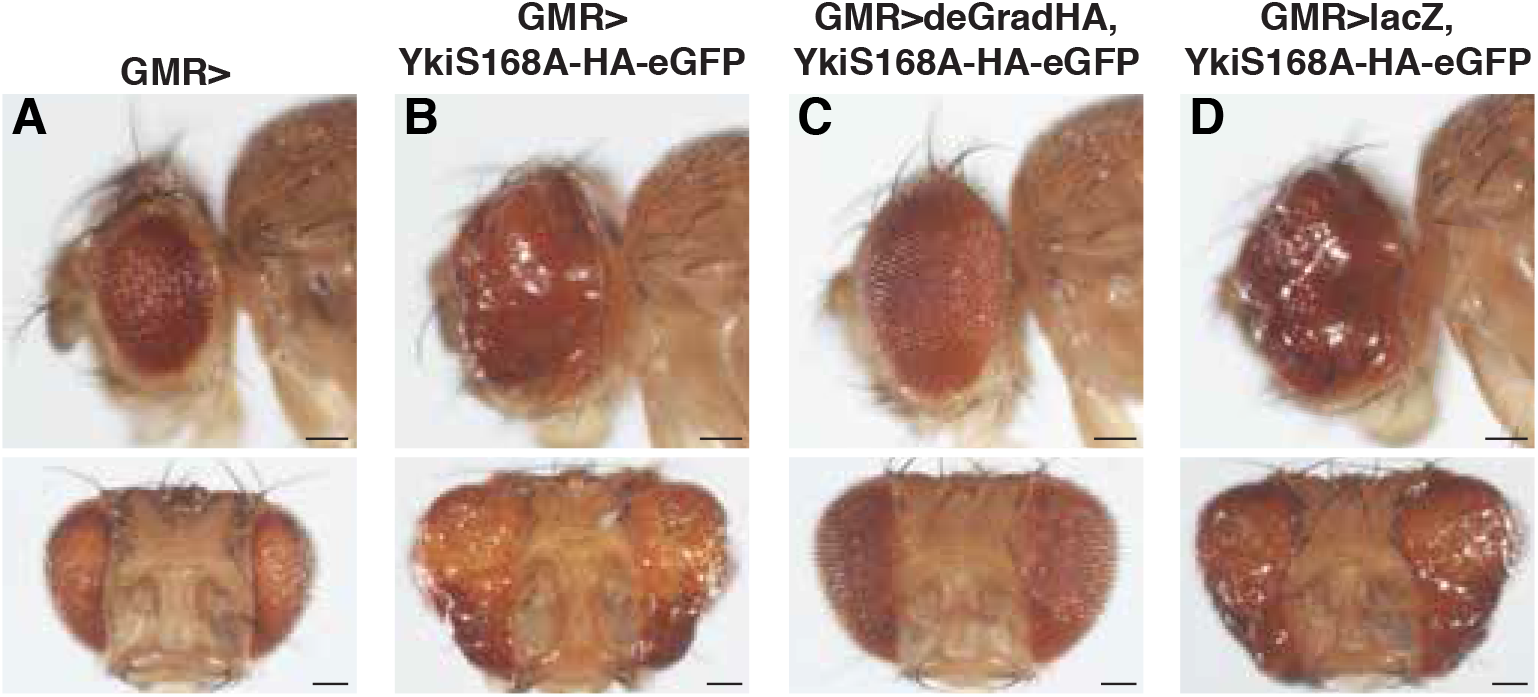
Functionalization of anti-HAscFv Frankenbody X15F11 (HA system) to degrade proteins *in vivo*. Side (top row) or frontal (bottom row) view of *Drosophila* adult eyes expressing the eye-specific GMR-GAL4 driver (**A**), YkiS168A-HA-eGFP under the control of GMR-GAL4 (**B**), YkiS168A-HA-eGFP as well as deGradHA under the control of GMR-GAL4 (**C**), or YkiS168A-HA-eGFP together with lacZ under the control of GMR-GAL4 (**D**). Scale bars are 100 µm.

## Discussion

### Single copies of short peptides and their binders

We focused our study on short peptide tags for which specific, high-affinity binders, which are soluble and functional in the intracellular milieu have been characterized. Therefore, we selected the following systems: SunTag (Tanenbaum et al., 2014), MoonTag (Boersma et al., 2019), HA (Zhao et al., 2019) and ALFA (Gotzke et al., 2019). For other commonly used tags such as FLAG® (Hopp et al., 1988) or Myc (Evan et al., 1985), we are unaware of specific binders derived from the corresponding monoclonal antibodies that could perform as intrabodies (Fujiwara et al., 2002; Marschall et al., 2015; Moutel et al., 2016; Worn et al., 2000).

Recently, a number of other short peptide binders were characterized, the BC2 nanobody recognizing the N-terminal aa 16-27 of beta catenin (Traenkle et al., 2015), the KTM219-derived scFv binding to a stretch of 7 aa of the BGPC7 (bone Gla protein or osteocalcin) (Wongso et al., 2017) and the nanobody NbSyn2 against the C-terminal of α-Synuclein (EPEA C-tag (De Genst et al., 2010)). Although they were shown to work intracellularly as chromobody or flashbody, we did not investigate them, since they recognize and bind to the corresponding endogenous proteins. Another binder, the nanobody VHH05 binding to a 14 aa peptide epitope of the E2 ubiquitin-conjugating enzyme UBC6e (Ling et al., 2019), was published after we had initiated our studies.

We also tested the OLLAS system (Park et al., 2008), since the OLLAS tag is synthetic and a high affinity monoclonal antibody was recently described and successfully used in several developmental studies (Nern et al., 2015; Wu et al., 2018; Yamazaki et al., 2016). Unfortunately, we were unable to demonstrate its possible utilization as an intracellular binder (using the scFv format) in the same experimental conditions (data not shown but available upon request).

We were particularly interested in testing whether the binders would be able to efficiently bind to a single copy of the selected tag *in vivo*. If this were the case, proteins of interest could be minimally modified with the aim not to affect any of their *in vivo* functions. Furthermore, current technology of precise gene knockin or tagging might be more efficient with short insertions in some organisms, such as zebrafish, for example. With the exception of the ALFA system and HA-frankenbody, the SunTag and MoonTag systems were previously tested *in vivo* in a similar setup to ours but with cellular baits containing multiple copies of the corresponding tag, as the primary interest of the authors was to visualize *in vivo* translation at a single molecule resolution.

### Influence of tags on the baits

We did observe that the various tags, even in single copy, could mildly alter the expression of the POI examined. The insertion of ALFA and MoonTag into the mitochondrial bait (mito_mCherry_MoonTggp41 and mito_mCherry_ALFA) slightly altered the mitochondrial “shape” and resulted in residual cytoplasmic signal upon overexpression (Suppl. Figure 1a). This may reflect a higher expression level of these fusions leading to higher background and/or slight disturbance of the mitochondria or, alternatively, could be a direct effect of the peptide tag. Please note that Götzke et al. (Gotzke et al., 2019) used a slightly different mitochondrial bait with one copy of ALFA tag and they did not report a similar pattern of expression; moreover, the same mitochondrial bait with 12 copies of the MoonTaggp41 was tested in another cellular context (Boersma et al., 2019).

The insertion of the HA tag into Vimentin (Vimentin_mCherry HA) also altered the appearance of the filaments painted by the mCherry signal (Suppl. Figure 1a); again, we think it is more likely a consequence of overexpression rather than a direct influence of the specific tags (or mCherry-tag(s) module).

All the nuclear baits localized exclusively to the nucleus, irrespective of the nature or position of the tags.

### Expression of the binders

We confirmed that all the tested small tag binders, the scFvs (anti-SunTag and anti-HA frankenbodies) and the nanobodies (anti-MoonTag and anti-ALFA), were excellent intrabodies and chromobodies; they were expressed at high levels inside the cells and diffused freely both in the cytoplasm and in the nucleoplasm. We occasionally observed some minor aggregation with the anti-GCN4scFv, probably due to the high overexpression from a CMV promoter, and with both anti-HA frankenbodies. Moreover, the nanobodies binding MoonTag and ALFA hardly displayed any aggregation when expressed at high level, confirming the high solubility of these protein binders (Beghein and Gettemans, 2017; Ingram et al., 2018; Schumacher et al., 2018).

The type of FP chosen for the generation of chromobodies (Kaiser et al., 2014; Keller et al., 2019; Moutel et al., 2018) may partially influence its expression and/or function: we noticed that for the anti-ALFA Nb, which was originally tested with mScarlet (Gotzke et al., 2019), fusion to sfGFP was preferable, since we observed a weak interference of the mEGFP over the mCherry signal of some baits; mVimentin_mCherry_ALFA, for example, had a lower intensity signal when bound to the anti-ALFANb_mEGFP than to the anti-ALFANb_sfGFP, or mVimentin_mCherry_HA when bound to both frankenbodies, which were also fused to mEGFP, than when expressed alone. However, fusion to mEGFP resulted in higher and brighter signals, especially in the nuclei.

Overall, we showed that all the binders tested were able to recognize and bind *in vivo* a single copy of the respective peptide tag embedded in proteins of different cell compartments, albeit with different efficiency and affinity.

The systems based on nanobodies (MoonTag and ALFA) might be more suitable for experiments in nuclear and subnuclear compartment, given their general higher “solubility” inside the cell.

We did not notice significant differences of the SunTag, MoonTag or ALFA for recruiting the respective binders to the mitochondria, to membranes or to filaments. The single HA tag, in our cellular experiments, was sufficient to bind and recruit the corresponding frankenbodies to all the structures analysed, but displayed a lower affinity than the three other tags, in agreement with the lower signal to noise ratio of the Mito_mCherry_1xHA versus Mito_mCherry_smHA in cells, or of 10xHA - H2B-mCherry versus 1x or 4xHA in zebrafish, reported by Zhao (Zhao et al., 2019). The lower binding affinity might also correlate with the size of epitope, as the HA tag is the smallest (9 amino acids) Furthermore, our experiments in *Drosophila* confirmed the positive correlation of the HA copy number and *in vivo* binding. While this could represent a drawback of the HA system, it might also provide an opportunity for titrating the effects of functionalized HA binders by adjusting the number of the HA copies fused to the POI.

### Combination of multiple tags

Combinatorial tagging of POIs would expand the repertoire of protein manipulation. A possibility would be, for example, to use one tag and its specific binder for visualization (with a chromobody) and the other tag for specific manipulation (see next paragraph). As pointed out in a recent review (Aguilar et al., 2019b), expression levels of the protein binder for visualization of a POI must be carefully controlled for a correct interpretation of the results. Strategies such as inducibility (Panza et al., 2015), self-transcriptional autoregulating domain fusion (Son et al., 2016) or intrinsic self stability (Tang et al., 2016), which were developed for nanobodies (Panza et al., 2015; Tang et al., 2016) and fibronectin-derived intrabodies (Son et al., 2016), could be applied to all the small tag binders described here.

Our control experiments using baits containing tags that were not supposed to be recognized by the different binders revealed some cross-reactivity between the SunTag and the ALFA systems, mostly with the anti-GCN4 scFv recognizing the ALFA tag rather than the reverse (Suppl. Figure 3, panels of G and J). The similarity of the two tags consists in a stretch of only 3 aa (EEL), but this might be sufficient for low affinity binding. No other significant cross-reactivity was observed among the other systems, confirming the suitable orthogonality described for SunTag and MoonTag by Boersma et al. (Boersma et al., 2019). Any combination of two or even three tags would certainly be beneficial for some experiments, with the avoidance of SunTag/ALFA pair.

Inserting several tags into an endogenously expressed protein will also allow for efficient recognition of the protein by an antibody against one of the tags, and manipulation of the POI via a functionalized binder recognizing the second tag inserted. The effects of the manipulation can then be followed with antibody staining (see (Aguilar et al., 2019b) for a discussion on the use of several different tags in the same gene).

### Functionalization of small tag binders

We demonstrate the ability of the binders to be recruited by “single-tagged” anchored proteins to different cellular compartment. The reverse approach, that is, move or trap the “single-tagged” POI with an anchored binder, is a possible functionalization of these small tag binders. Mislocalization or trapping of some POIs, tagged with FP, has been developed with anti-GFP nanobodies (Harmansa et al., 2017; Seller et al., 2019), anti-mTFP DARPin (Vigano et al., 2018), anti-mCherry nanobody (Prole and Taylor, 2019) but also with nanobodies against endogenous, non-tagged proteins, for example Gelsolin and CapG (Van Audenhove et al., 2013).

Another functionalization that can be applied to the small tag binders is the addition of “degrons” (Natsume and Kanemaki, 2017) to achieve specific and temporally-controlled degradation of the tagged POIs. This approach has been successfully applied using the anti-FP nanobodies (Aguilar et al., 2019a; Beghein and Gettemans, 2017; Deng et al., 2020; Ingram et al., 2018; Prole and Taylor, 2019). Here we expand these findings by demonstrating that a single short HA tag can be efficient applied to inactivate POIs using deGradHA, a tool adapted from the deGradFP system and designed to channel HA-tagged POIs to ubiquitin/proteasome dependent degradation. Thus, in addition to relocalization, small tags in single copies can be used to target POIs for proteolysis enabling a spectrum of additional applications.

Finally, addition of any enzymatic domain to the the small tag binders would specifically modify the tagged POI, as it was elegantly shown with a minimal Rho kinase domain fused to the GFP nanobody to phosphorylate a GFP tagged protein in *Drosophila melanogaster* (Roubinet et al., 2017) or a proximity-directed O- GlcNAcetylation by linking the O-GlcNAc transferase activity to the GFP or EPEA nanobody in cell culture (Ramirez et al., 2020). Several recent reviews highlighted the versatility of the nanobodies for numerous applications both in clinical and biological research (Beghein and Gettemans, 2017; Ingram et al., 2018; Schumacher et al., 2018). The various functionalization strategies can be extended to these small tag binders.

An important aspect in developing tools and strategies for acute protein manipulation in cultured cells and in living organisms is the temporal and spatial inducibility and/or reversibility of the manipulation itself. Recent publications have demonstrated the possibility to directly modify certain nanobodies in order to control their binding to the target protein either with light (Gil et al., 2019; Yu et al., 2019) or with small molecules (Farrants et al., 2020). It will be exciting to extend these types of modification to the small tag binders in order to achieve this extra level of regulation and expand the toolbox to acutely and reversibly manipulate proteins *in vivo*.

## MATERIALS and METHODS

### Plasmid construction

All the eukaryotic expression plasmids were generated by specific PCR amplification and standard restriction cloning. Briefly, the mitochondrial baits containing an N-terminal anchor sequence from the human CISD1 protein (the first 59 amino acids) fused to the N-terminus of mTFP1, were generated from pcDNA4TO-mito-mCherry-10xGCN4_v4 (Addgene plasmid 60914 (Tanenbaum et al., 2014)) by substituting the 10xGCN4_v4 with each individual tag, PCR amplified with specific primers and inserted with RsrII/SacII sites.The pH2B_mCherry_Tag plasmids were generated from the respective pmito_mCherry_Tag, substituting the CISD1 protein with the human H2BC11 (Histone H2B) by restriction cloning. The other nuclear baits pTag_mCherry_H2b and pmCherry_Tag_H2B were also generated by inserting each PCR amplified Tag into pmCherry_H2B (a kind gift from E.Nigg group). Substitution of CISD1 with PCR amplified mouse Vimentin inserted at EcoRI/BamHI sites of each pmito_mCherry_Tag generated the filaments baits. ptwistmCherry_CD8_OLLAS_SunTagv4 was synthetized at TWIST® Bioscience (South San Francisco, CA).

The anti-GCN4-scFv was generated from pHR-scFv-GCN4-sfGFP-GB1-dWPRE (Addgene plasmid 60907 (Tanenbaum et al., 2014), cut with EcoRI/XbaI and inserted into pcDNA3. The anti-gp41Nb was generated from pHR-Nb 2H10 gp41- sfGFP-GB1-dWPRE (a kind gift from the M.Tannenbaum group)(Boersma et al., 2019), cut with EcoRI/XbaI and inserted into pcDNA3. The anti-HAscFvs frankenbodies were kindly provided by T.Stasevich (Zhao et al., 2019). For the anti-ALFA nanobody (Gotzke et al., 2019) either sfGFP-GB1 or mEGFP were PCR amplified and inserted at the BamH1/Not1 site of pNT-NAM01 pCMV-NbALFA- MCS, kindly provided by S.Frey.

For *Drosophila* expression pUASTLOTattB_frankenbody_aHA- scFvX15F11_mEGFP was generated by cutting the frankenbody_aHA- scFvX15F11_mEGFP with XhoI/XbaI and inserting the frankenbody into pUASTLOTattB (Kanca et al., 2014). For pUASTLOTattB_deGradHA, vhhGFP4 of pUAST_NSlmb-vhhGFP4 (Addgene plasmid 35575 (Caussinus et al., 2013)) was cut out and replaced with frankenbody_aHA-scFvX15F11 amplified by PCR. The resulting plasmid was cut with EcoRI/XbaI to insert deGradHA into pUASTLOTattB. All constructs were verified by sequencing. Plasmid maps and oligonucleotide sequences for PCR and cloning are available upon request. A schematic representation of the fusion constructs is provided in Figure 1

### Cell cultures, transfections and imaging

HeLa S3α cells, routinely tested for mycoplasm contamination, were maintained in Dulbecco’s modified Eagle’s medium supplemented with 10% foetal calf serum, 100 IU penicillin and 100 μg streptomycin per ml. One day before transfection, cells were seeded on glass cover slips placed into a 24 well plate at a density of 50,000-100,000 cells/well.

Transfections were carried out with 1 μg of total DNA (500 ng for each construct or with empty expression plasmid) and 3 μl of FuGENE ® HD Transfection Reagent (Promega), according to the manufacturer’s instructions. 24 hours post transfection, cells were fixed in 4% paraformaldehyde, stained with Hoechst 33342 (Invitrogen) and mounted on standard microscope slides with VECTASHIELD® (Vector Laboratories Inc. Burlingame, CA).

Confocal images were acquired with a Leica point scanning confocal “SP5-II-MATRIX” microscope (Imaging Core Facility, Biozentrum, University of Basel) with a 63x HCX PLAN APO lambda blue objective and 1-2x zoom.

### Drosophila lines

Transgenic *Drosophila* lines carrying a UASTLOT_frankenbody_aHA- scFvX15F11_mEGFP or UASTLOT_deGradHA insertion in chromosomal position Chr3L, 68A4 (attP2) were generated by standard procedures using PhiC31/attB- mediated integration.

UASpH2Av::Flag-HA (H2Av-Flag-1xHA) flies were a kind gift of the N. Iovino group (Max Planck Institute for Immunobiology and Epigenetics, Freiburg, Germany). UASHistone4-3xHA (H4-3xHA) flies were created by the FlyORF Zurich ORFeome Project (Bischof et al., 2013) (Fly Line ID F000777). Brk-GAL4 (53707), GMR-GAL4 (1104), UASYkiS168A-HA-eGFP (28836; described in (Oh and Irvine, 2008)) and UASlacZ (28836) flies were provided by the Bloomington *Drosophila* Stock Center.

### Immunohistochemistry and imaging of Drosophila samples

Salivary glands from third instar *Drosophila* larvae were dissected, fixed and stained using standard procedures. The following antibodies were used: rabbit anti-GFP (1:500, Abcam), rat anti-HA (1:200, Roche), mouse anti-Flag (1:500, Sigma), Alexa fluorophore-conjugated secondary antibodies (1:500; A11034, A11031, A11077) and Hoechst 33342 (1:5000; Invitrogen). Images were acquired using a Zeiss LSM880 laser scanning confocal microscope (Life Imaging Center (LIC), Center for Biological Systems Analysis (ZBSA), Albert-Ludwigs University Freiburg).

## Acknowledgments

We would like to thank Simon Ittig, Christoph Kasper, Marilise Amstutz of T3 Pharmaceuticals for their generosity to host one of us (MAV). We are indebted to Dafne Iberra-Morales and Nicola Iovino (Max Planck Institute of Immunobiology and Epigenetics, Freiburg, Germany) for the H2Av-Flag-1xHA flies. We thank the staff of the Life Imaging Center (LIC) in the Center for Biological Systems Analysis (ZBSA) of the Albert-Ludwigs-University Freiburg for confocal microscopy resources and support in image recording. We also thank the Imaging core facility of the Biozentrum for their assistance, all the members of Affolter lab for helpful discussions and Bernadette Bruno, Gina Evora and Karin Mauro for their great help in the media kitchen.

The work in the Affolter lab was supported in parts by grants from SNF and SystemsX (MorphogenetiX) as well as by the Kantons Basel-Stadt and Basel-Land. Work in the Pyrowolakis lab was funded by the Deutsche Forschungsgemeinschaft (DFG, German Research Foundation) under Germany s Excellence Strategy EXC- 2189-Project ID 390939984.

## FIGURE Legends

Supplementary Figure 1a

**Intracellular expression of mitochondrial, filament and membrane baits**

Confocal images of HeLa cells transiently transfected with **(A)** pmito_mCherry_SunTagV4, **(B)** pmito_mCherry_MoonTaggp41, **(C)** pmito_mCherry_HA, **(D)** pmito_mCherry_ALFA, **(E)** pmVimentin_mCherry_SunTagV4, (**F**) pmVimentin_mCherry_MoonTaggp41, (**G**) pmVimentin_mCherry_HA, (**H**) pmVimentin_mCherry_ALFA, (**I**) ptwist_mCherry_CD8_OLLAS_SunTagV4. The first column of each row indicated by the letter represents the mCherry channel (red), the second column is the nuclear Hoechst staining (blue) and the third column is the overlay of the two channels channels (with the scale bar in white (15 μm)), showing the localization of the mitochondrial (**A-D**), filaments (**E-H**) and membrane (**I**) baits; Images were taken 24 hours post transfection. Transfected constructs are indicated at bottom of each row and the single and merge channels are indicated at the top of the respective columns. The figures are from a representative experiment, performed at least three times.

Supplementary Figure 1b

**Intracellular expression of the nuclear baits**

Confocal images of HeLa cells transiently transfected with **(A)** pH2B_mCherry_SunTagV4, **(B)** pH2B_mCherry_MoonTaggp41, **(C)** pH2B_mCherry_HA, **(D)** pH2B_mCherry_ALFA, **(E)** pmCherry_SunTagV4_H2B, (**F**) pmCherry_MoonTaggp41_H2B, (**G**) pmCherry_HA_H2B, (**H**) pmCherry_ALFA_H2B, (**I**) pSunTagV4_mCherry_H2B, (**J**) pMoonTaggp41_mCherry_H2B, (**K**) pHA_mCherry_H2B, (**L**) pALFA_mCherry_H2B, (**M**) pmCherry_H2B. The first column of each row indicated by the letter represents the mCherry channel (red), the second column is the nuclear Hoechst staining (blue) and the third column is the overlay of the two channels channels (with the scale bar in white (15 μm)), showing the localization of the nuclear baits; Images were taken 24 hours post transfection. Transfected constructs are indicated at bottom of each row and the single and merge channels are indicated at the top of the respective columns. The figures are from a representative experiment, performed at least three times.

Supplementary Figure 2

**Intracellular binding of anti-GCN4 scFv and anti-gp41 Nanobody to nuclear baits**

Confocal images of HeLa cells transiently transfected with **(A)** pcDNA_aGCN4- scFv_sfGFP_GB1 alone; the combination of pcDNA_aGCN4-scFv_sfGFP_GB1 and **(B)** pSunTagv4_mCherry_H2B; **(C)** pH2B_mCherry_SunTagv4; **(D)** pcDNA_agp41- Nb2H10_sfGFP_GB1 alone; the combination of pcDNA_ agp41- Nb2H10_sfGFP_GB1 and **(E)** pmCherry_MoonTaggp41_H2B; **(F)** pH2B_mCherry_MoonTaggp41. The first column represents the GFP channel (green), the second column is the mCherry channel (red), the third column is the overlay of the two channels, showing the colocalization (indicated in yellow) of the antiGCN4 scFvs (**A-C**) or the anti-gp41Nb (**D-F**) with the respectively tagged nuclear baits; the fourth column represents the nuclear Hoechst staining (blue) and the fifth column is the merge of all three channels (with the scale bar in white (15 μm) on the bottom right corner). Images were taken 24 hours post transfection. Transfected constructs are indicated at the left of each row and the single and merge channels are indicated at the top of the respective columns. The figures are from a representative experiment, performed at least three times.

Supplementary Figure 3

**Negative controls of anti-GCN4 scFv**

Confocal images of HeLa cells transiently transfected with the combination of pcDNA_aGCN4-scFv_sfGFP_GB1 and **(A)** pmito_mCherry_HA, (**B**) pmito_mCherry_MoonTaggp41, **(C)** pHA_mCherry_H2B, **(D)** pmCherry_HA_H2B, **(E)** pH2B_mCherry_HA, (**F**) pH2B_mCherry_MoonTaggp41, (**G**) pH2B_mCherry_ALFA, (**H**) pmCherry_H2B, (**I**) pmVimentin_mCherry-MoonTaggp41, (**J**) pmVimentin_mCherry_ALFA. The first column represents the GFP channel (green), the second column is the mCherry channel (red), the third column is the overlay of the two channels, showing the colocalization (indicated in yellow) of the antiGCN4 scFv with mitochondrial (**A-B**), nuclear (**C-H**),) and filaments (**I-J**) baits carrying different tags; the fourth column represents the nuclear Hoechst staining (blue) and the fifth column is the merge of all three channels (with the scale bar in white (15 μm) on the bottom right corner). Images were taken 24 hours post transfection. Transfected constructs are indicated at the left of each row and the single and merge channels are indicated at the top of the respective columns. The figures are from a representative experiment.

Supplementary Figure 4

**Negative controls of anti-gp41 Nanobody**

Confocal images of HeLa cells transiently transfected with the combination of pcDNA_ agp41-Nb2H10_sfGFP_GB1and **(A)** pmito_mCherry_HA, (**B**) pmito_mCherry_SunTagv4, **(C)** pmito_mCherry_ALFA **(D)** pH2B_mCherry_ALFA, **(E)** pH2B_mCherry_SunTagv4, (**F)** pmCherry_H2B, (**G**) pmVimentin_mCherry-SunTagv4, (**H**) pmVimentin_mCherry_ALFA. The first column represents the GFP channel (green), the second column is the mCherry channel (red), the third column is the overlay of the two channels, showing the colocalization (indicated in yellow) of the antigp41-Nb with mitochondrial (**A-C**), nuclear (**D-F**), and filaments (**G-H**) baits carrying different tags; the fourth column represents the nuclear Hoechst staining (blue) and the fifth column is the merge of all three channels (with the scale bar in white (15 μm) on the bottom right corner). Images were taken 24 hours post transfection. Transfected constructs are indicated at the left of each row and the single and merge channels are indicated at the top of the respective columns. The figures are from a representative experiment.

Supplementary Figure 5

**Intracellular binding of anti-HAscFv frankenbody X2E2 and extra nuclear coloclization of anti_HA frankenbody X15F11 (HA system)**

Confocal images of HeLa cells transiently transfected with **(A)** p_frankenbody_aHA- scFvX2E2_mEGFP alone; the combination of p_frankenbody_aHA- scFvX2E2_mEGFP and **(B)** pmito_mCherry_HA; **(C)** pHA_mCherry_H2B; **(D)** pmCherry_HA_H2B; (**E**) pH2B_mCherry_H2B; (**F**) pmVimentin_mCherry_HA. The confocal images in lower black frame represent the cotransfection of p_frankenbody_aHA-scFvX15F11_mEGFP with (**A’**) pHA_mCherry_H2B or (**B’**) pH2B_mCherry_HA.The first column represents the GFP channel (green), the second column is the mCherry channel (red), the third column is the overlay of the two channels, showing the colocalization (indicated in yellow) of the antiHAscFvs with the respective mitochondrial (**B**), nuclear (**C-E, A’-B’**) and filaments (**F**) baits; the fourth column represents the nuclear Hoechst staining (blue) and the fifth column is the merge of all three channels (with the scale bar in white (15 μm) on the bottom right corner). Images were taken 24 hours post transfection. Transfected constructs are indicated at the left of each row and the single and merge channels are indicated at the top of the respective columns. The figures are from a representative experiment, performed at least three times.

Supplementary Figure 6

**Negative mitochondrial and membrane controls of anti-HA scFvs**

Confocal images of HeLa cells transiently transfected with the combination of pfrankenbody_aHA-scFvX15F11_mEGFP (**A,C,E**) or pfrankenbody_aHA- scFvX2E2-mEGFP (**B,D**) and **(A-B)** pmito_mCherry_SunTagv4, (**C-D**) pmito_mCherry_MoonTaggp41, (**E**) ptwist_mCherry_CD8_OLLAS_SunTagv4. The first column represents the GFP channel (green), the second column is the mCherry channel (red), the third column is the overlay of the two channels, showing the colocalization (indicated in yellow) of the anti HA-scFvs with mitochondrial (**A-D**), and membrane (**E**) baits carrying different tags; the fourth column represents the nuclear Hoechst staining (blue) and the fifth column is the merge of all three channels (with the scale bar in white (15 μm) on the bottom right corner). Images were taken 24 hours post transfection. Transfected constructs are indicated at the left of each row and the single and merge channels are indicated at the top of the respective columns. The figures are from a representative experiment.

Supplementary Figure 7

**Negative nuclear and filaments controls of anti-HA scFvs**

Confocal images of HeLa cells transiently transfected with the combination of pfrankenbody_aHA-scFvX15F11_mEGFP (**A,C,E**) or pfrankenbody_aHA- scFvX2E2-mEGFP (**B,D,F**) and **(A-B)** pmCherry_H2B, (**C-D**) pH2B_mCherry_ALFA, **(E-F)** pmVimentin_mCherry_ALFA. The first column represents the GFP channel (green), the second column is the mCherry channel (red), the third column is the overlay of the two channels, showing the colocalization (indicated in yellow) of the anti HA-scFvs with nuclear (**A-D**), and filaments (**E-F**) baits carrying different tags; the fourth column represents the nuclear Hoechst staining (blue) and the fifth column is the merge of all three channels (with the scale bar in white (15 μm) on the bottom right corner). Images were taken 24 hours post transfection. Transfected constructs are indicated at the left of each row and the single and merge channels are indicated at the top of the respective columns. The figures are from a representative experiment.

Supplementary Figure 8

**Intracellular binding of anti-ALFA-Nb_mEGFP and extra nuclear colocalization of anti-ALFA-Nb_sfGFP_GB1**

Confocal images of HeLa cells transiently transfected with **(A)** pCMV_aALFA- Nb_mEGFP alone; the combination of pCMV_aALFA-Nb_mEGFP and **(B)** pmito_mCherry_ALFA; **(C)** pALFA_mCherry_H2B; **(D)** pmCherry_ALFA_H2B; (**E**) pH2B_mCherry_ALFA; (**F**) pmVimentin_mCherry_ALFA. The confocal images in lower black frame represent the cotransfection of pCMV_aALFA-Nb_sfGFP_GB1 with (**A’**) pHA_mCherry_ALFA_H2B or (**B’**) pH2B_mCherry_ALFA.The first column represents the GFP channel (green), the second column is the mCherry channel (red), the third column is the overlay of the two channels, showing the colocalization (indicated in yellow) of the antiALFA Nanobodies with the respective mitochondrial (**B**), nuclear (**C-E, A’-B’**) and filaments (**F**) baits; the fourth column represents the nuclear Hoechst staining (blue) and the fifth column is the merge of all three channels (with the scale bar in white (15 μm) on the bottom right corner). Images were taken 24 hours post transfection. Transfected constructs are indicated at the left of each row and the single and merge channels are indicated at the top of the respective columns. The figures are from a representative experiment, performed at least three times.

Supplementary Figure 9a

**Negative nuclear controls of anti-ALFA nanobodies**

Confocal images of HeLa cells transiently transfected with pCMV_aALFA- Nb_sfGFP_GB1 (**A,C,E** and **G**) or pCMV_aALFA-Nb_mEGFP (B,D,F and **H**)_alone **(A-B)** or in combination with (**C-D**) pH2B_mCherry_MoonTaggp41, **(E-F)** pH2B_mCherry_SunTagv4, (**G-H**) pmCherry_H2B. The first column represents the GFP channel (green), the second column is the mCherry channel (red), the third column is the overlay of the two channels, showing the colocalization (indicated in yellow) of the anti ALFA nanobody with nuclear (**C-H**) baits carrying different tags; the fourth column represents the nuclear Hoechst staining (blue) and the fifth column is the merge of all three channels (with the scale bar in white (15 μm) on the bottom right corner). Images were taken 24 hours post transfection. Transfected constructs are indicated at the left of each row and the single and merge channels are indicated at the top of the respective columns. The figures are from a representative experiment.

Supplementary Figure 9b

**Negative mitochondrial and filaments controls of anti-ALFA nanobodies** Confocal images of HeLa cells transiently transfected with the combination of pCMV_aALFA-Nb_sfGFP_GB1 (**A,C,E**) or pCMV_aALFA-Nb_mEGFP (B,D,F) and **(A-B)** pmito_mCherry_MoonTaggp41, (**C-D**) pmVimentin_mCherry_SunTagv4, **(E-F)** pmVimentin_mCherry_MoonTaggp41. The first column represents the GFP channel (green), the second column is the mCherry channel (red), the third column is the overlay of the two channels, showing the colocalization (indicated in yellow) of the anti ALFA Nanobodies with mitochondrial (**A-B**), and filaments (**C-F**) baits carrying different tags; the fourth column represents the nuclear Hoechst staining (blue) and the fifth column is the merge of all three channels (with the scale bar in white (15 μm) on the bottom right corner). Images were taken 24 hours post transfection. Transfected constructs are indicated at the left of each row and the single and merge channels are indicated at the top of the respective columns. The figures are from a representative experiment.

Supplementary Figure 10

**Intracellular binding of anti-HAscFv Frankenbody X15F11 (HA system) *in vivo*** Confocal images of *Drosophila* larval salivary glands expressing frankenbody_aHA- scFvX15F11_mEGFP alone (**A**), the nuclear bait H4-3xHA alone (**B**), or a combination of frankenbody_aHA-scFvX15F11_mEGFP and H4-3xHA (**C**). The first column represents the GFP channel (green, **A** and **C**) or the anti-HA staining channel (red, **B**). The second column represents the nuclear Hoechst staining (blue) and the third column is the merge of the two respective channels. Scale bars are 50 µm. Salivary glands were obtained from third instar *Drosophila* larvae expressing the constructs indicated at the left of each row. Single and merged channels are indicated at the top of the respective channel.

