## Supplementary Figures for "Protein manipulation using single copies of short peptide tags in cultured cells and in *Drosophila melanogaster*"

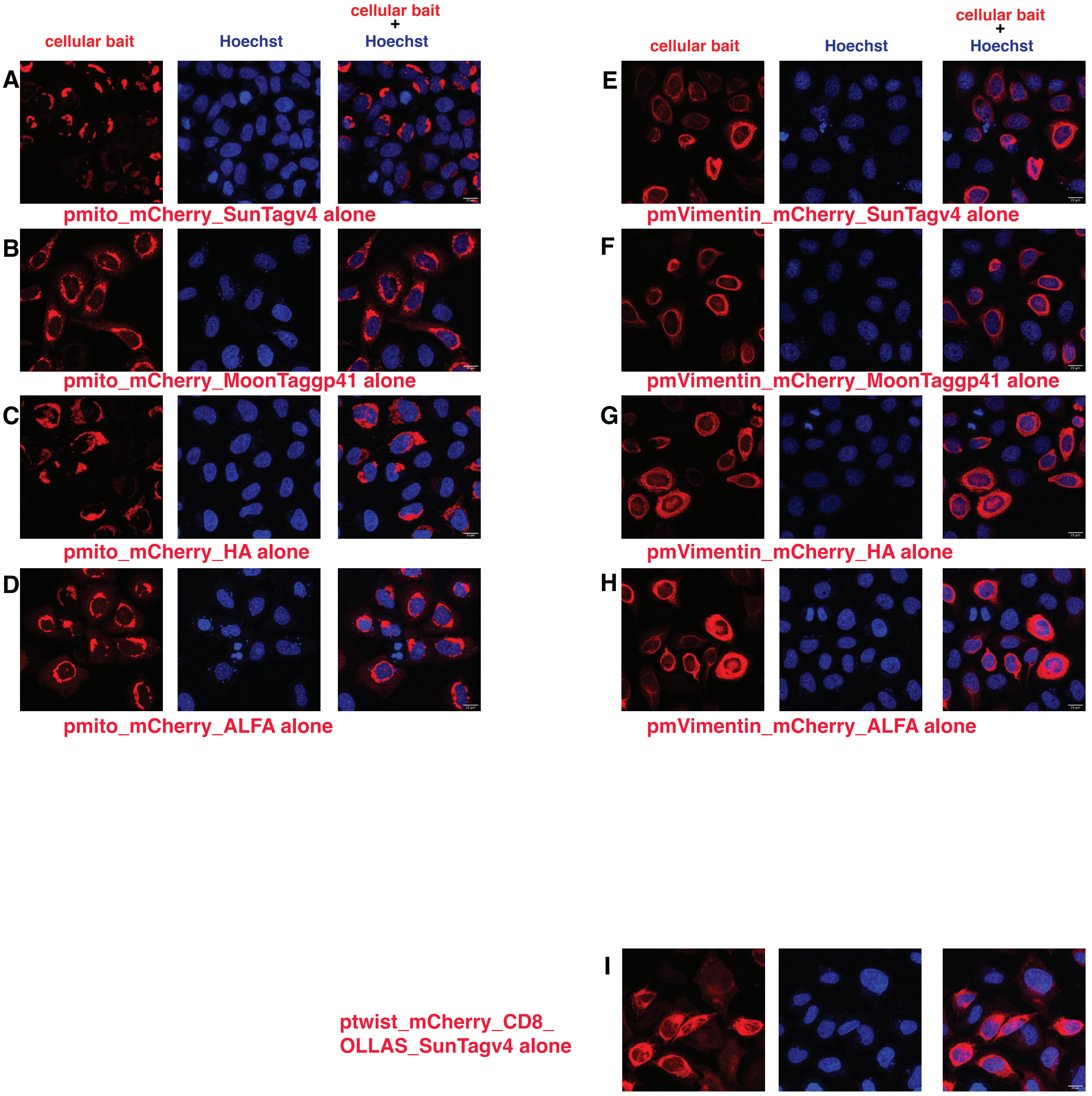


Vigano et al, Suppl. Figure1a

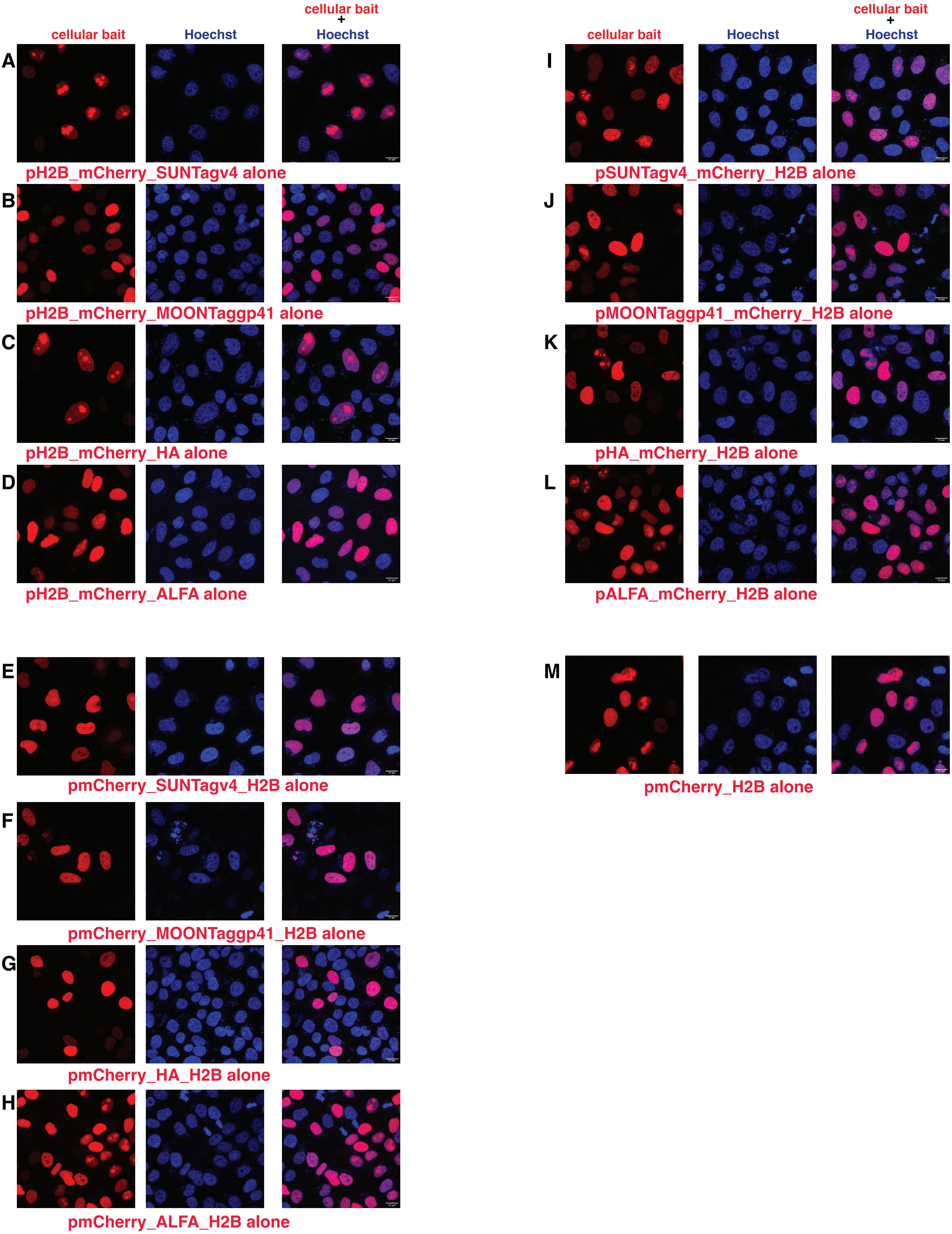


Vigano et al, Suppl. Figure1b

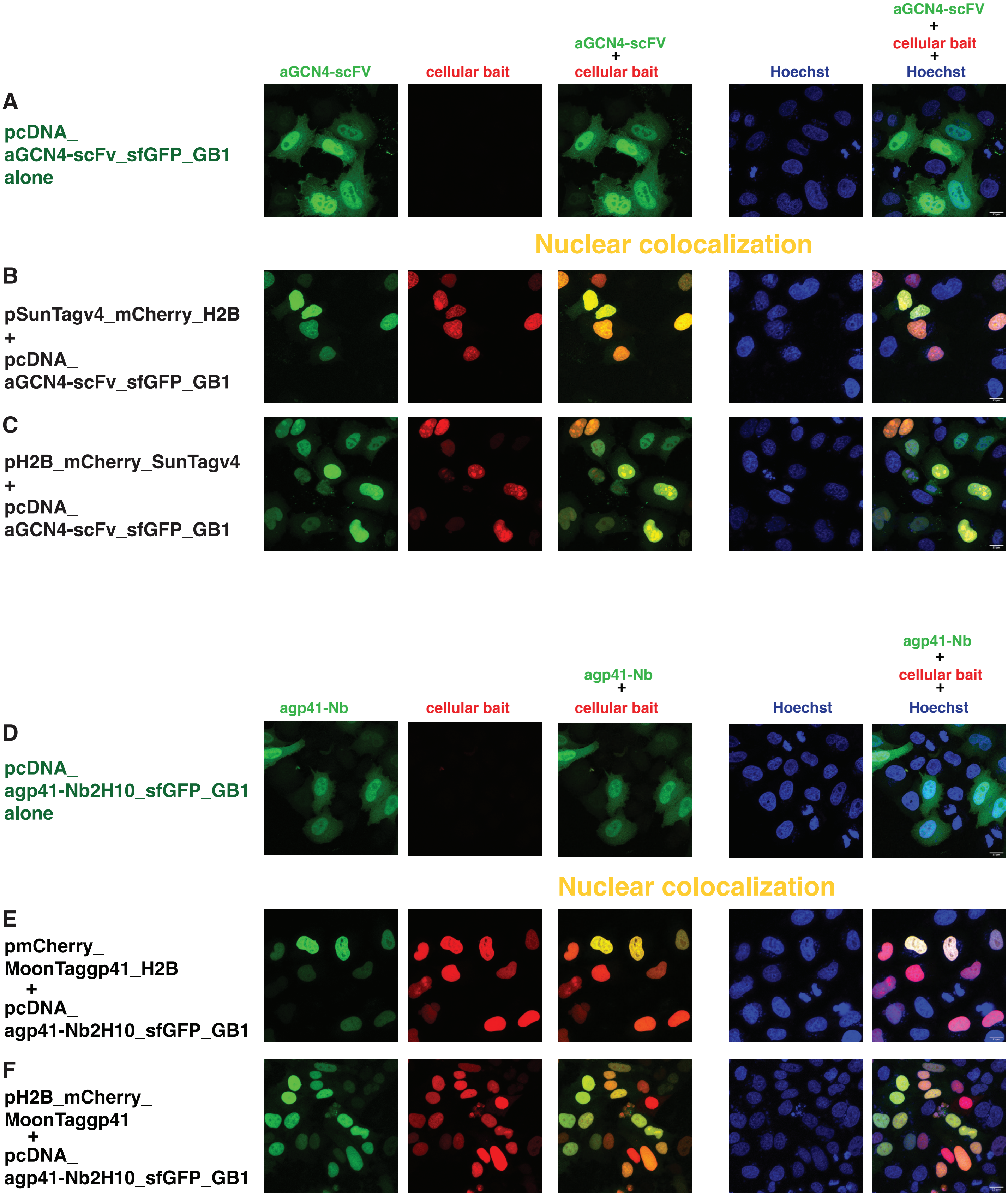


Vigano et al, Suppl. Figure2

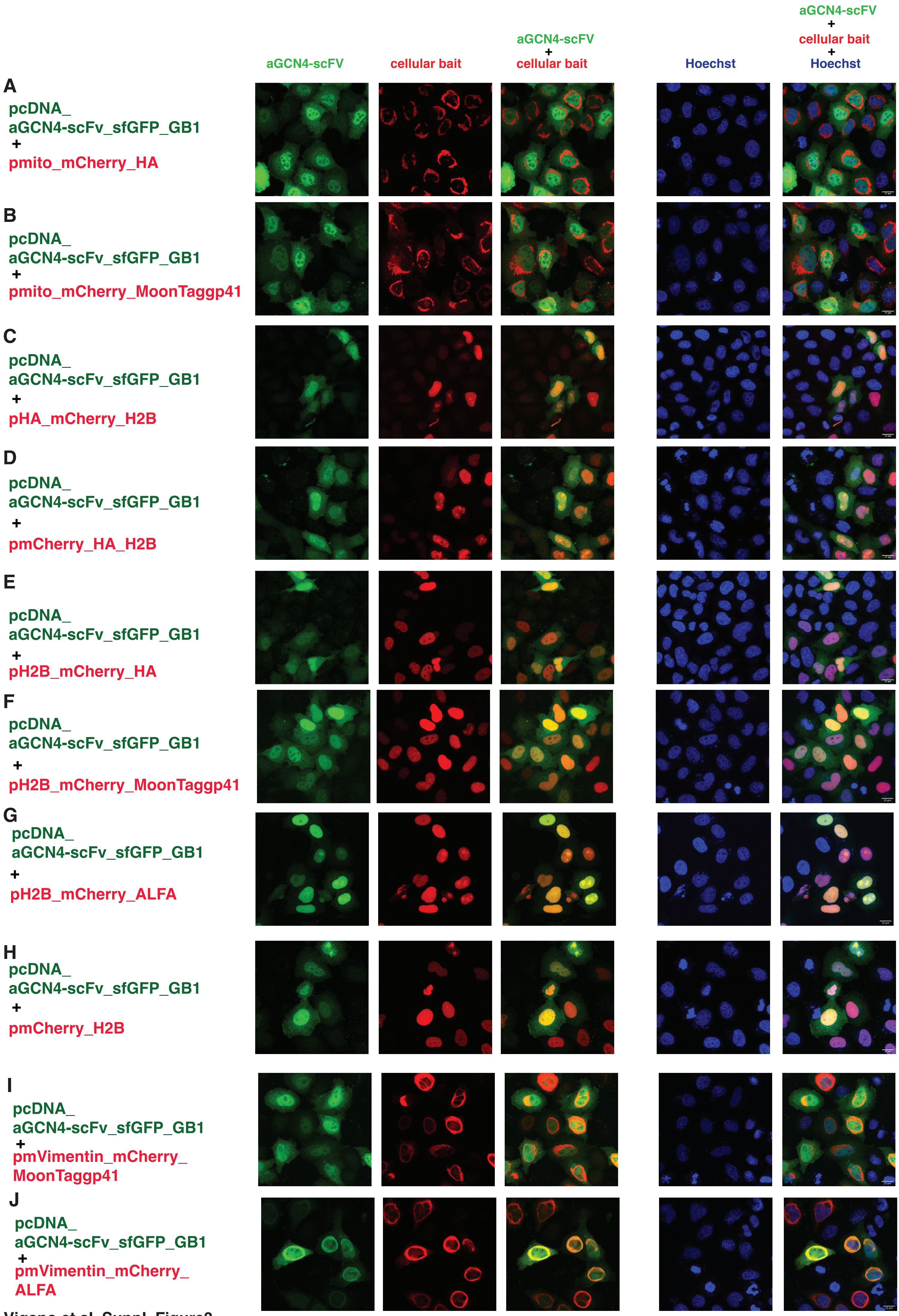


Vigano et al, Suppl. Figure3

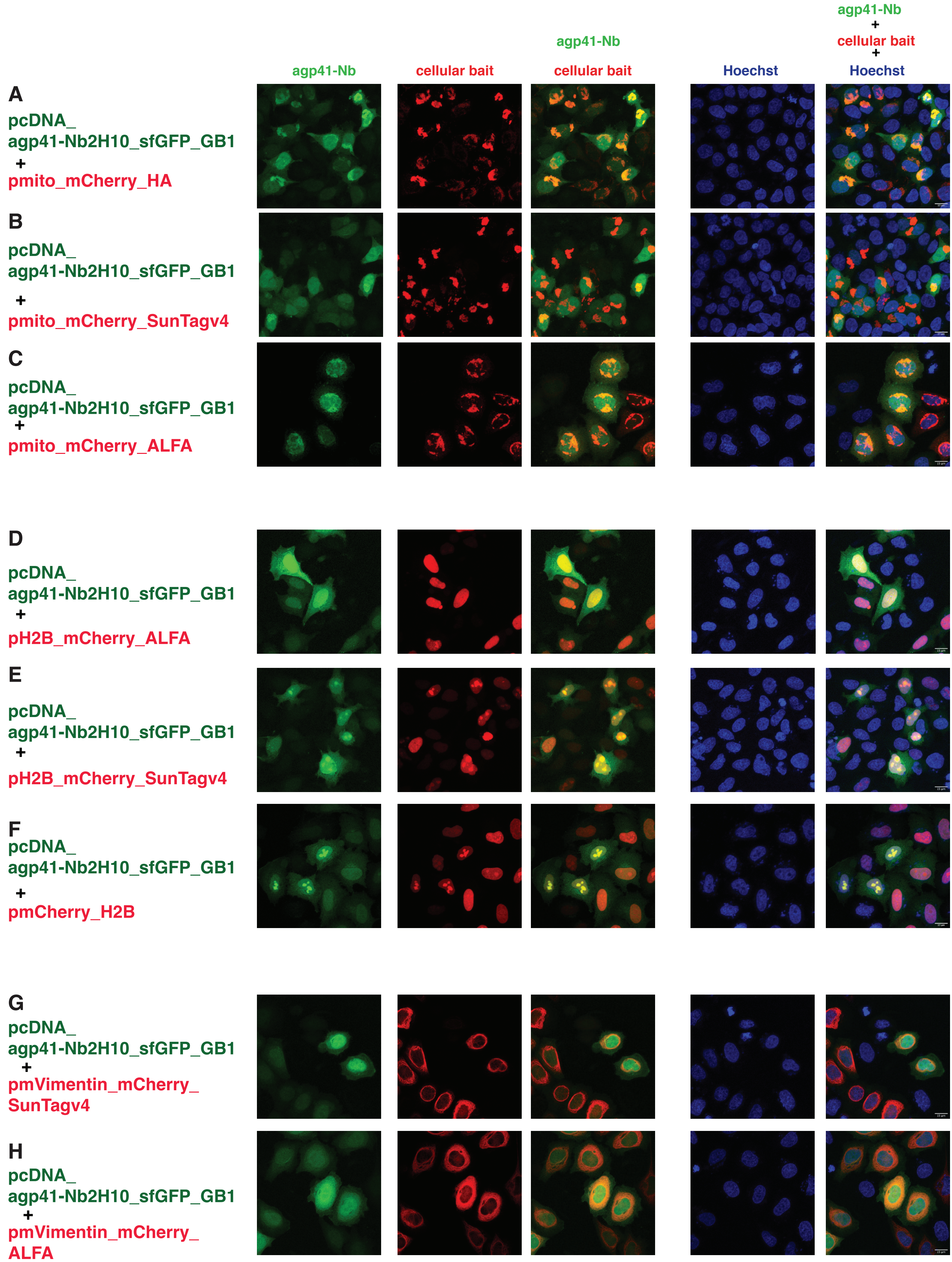


Vigano et al, Suppl. Figure4

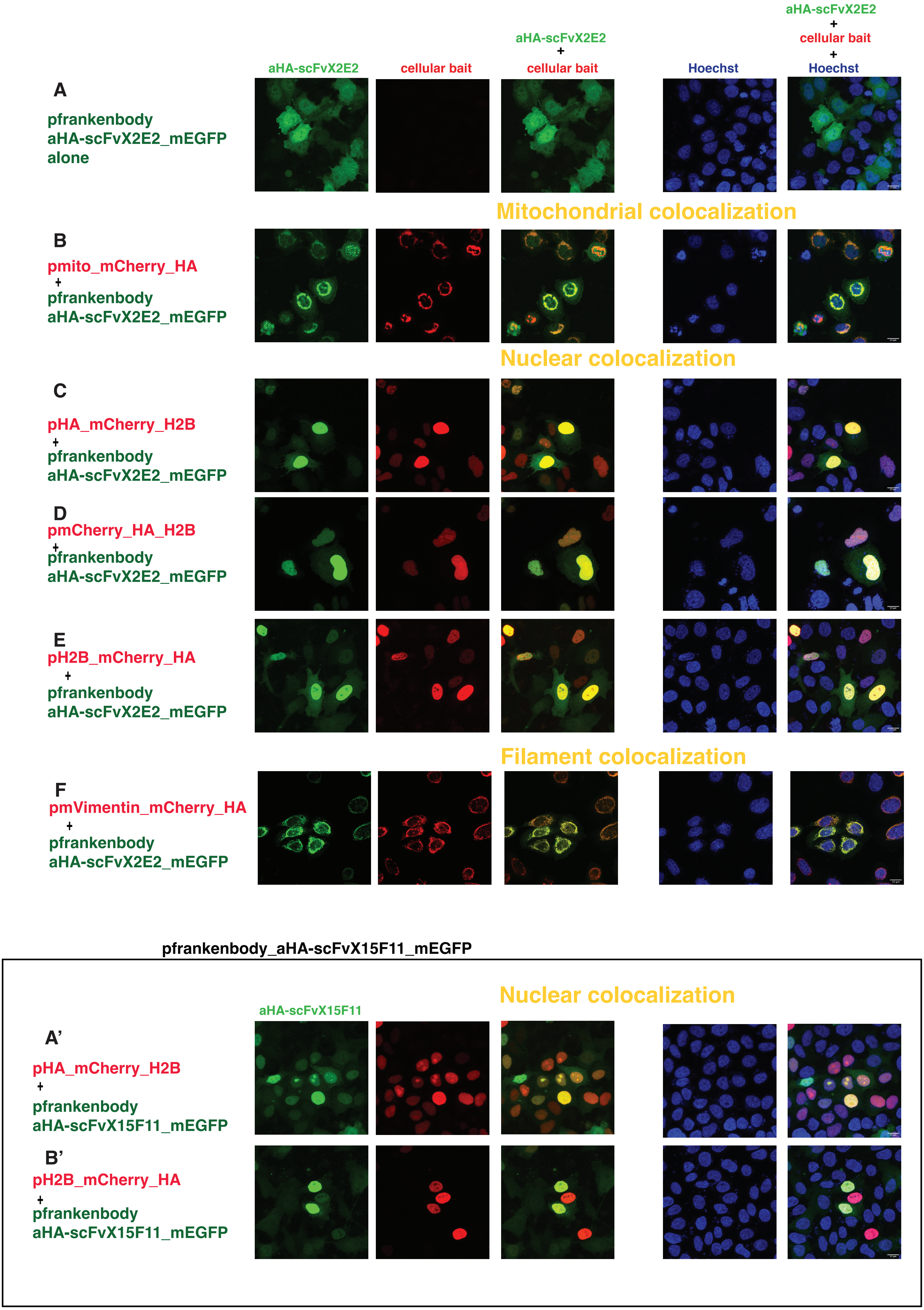


Vigano et al, Suppl. Figure5

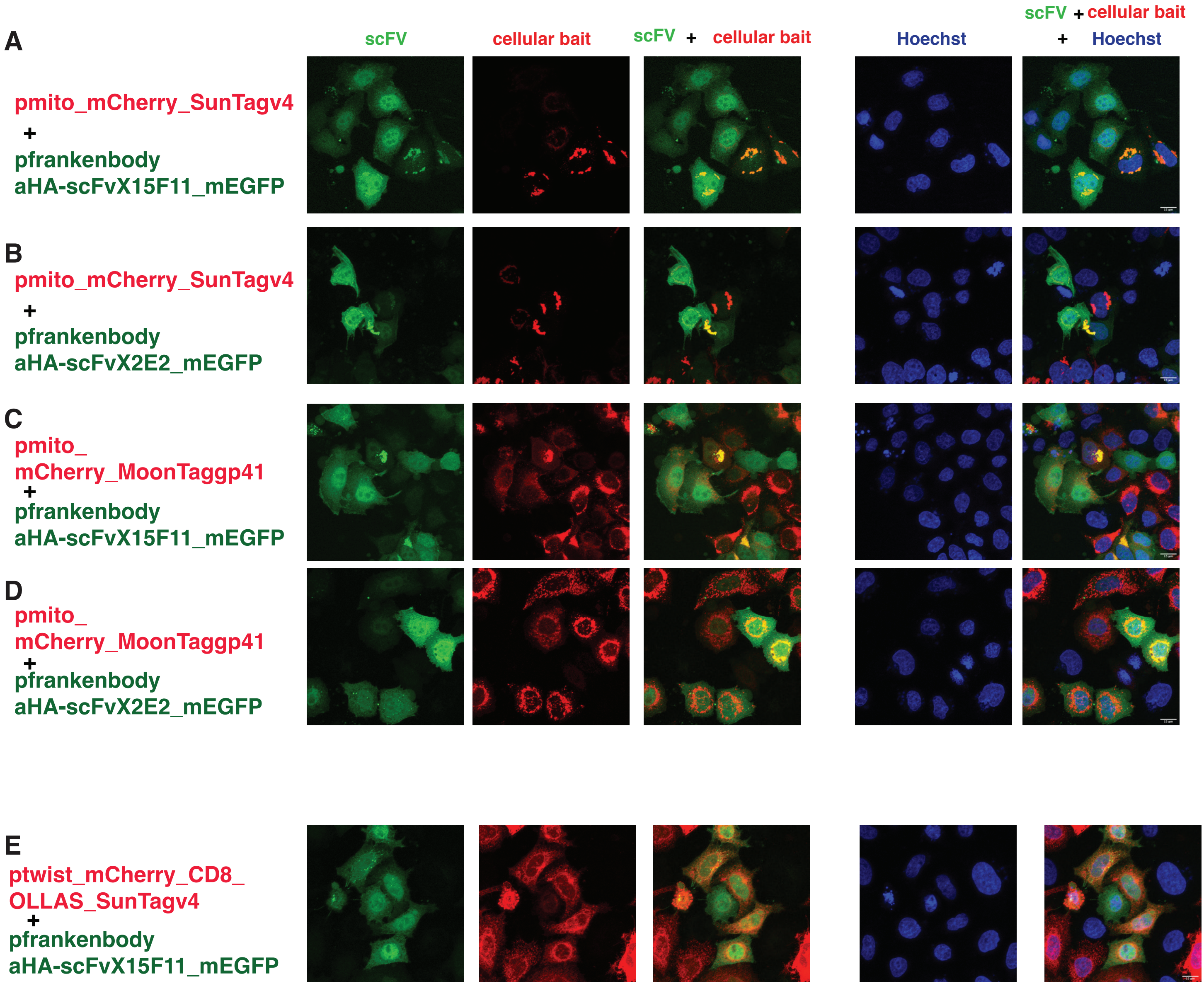


Vigano et al, Suppl. Figure6

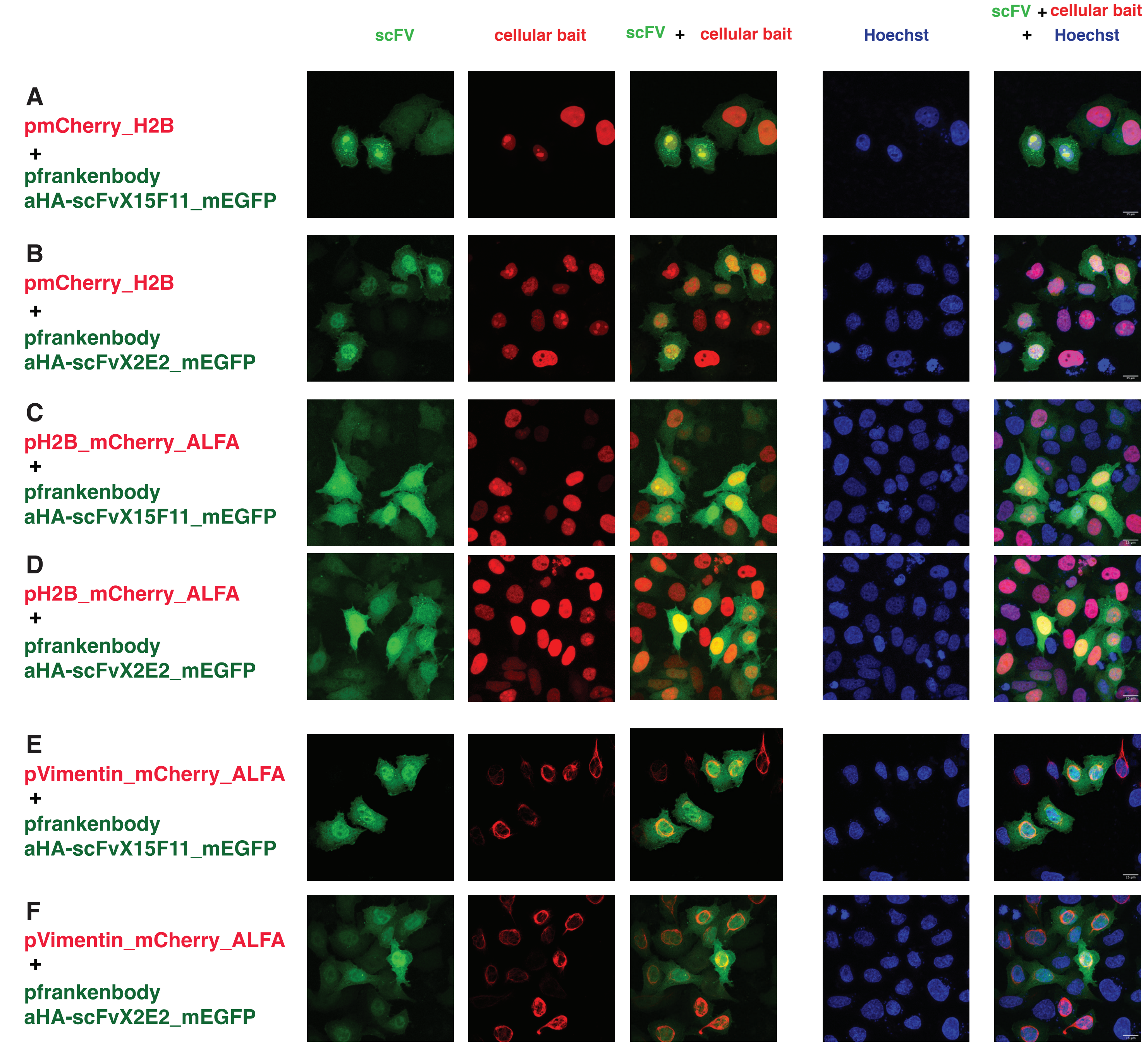


Vigano et al, Suppl. Figure7

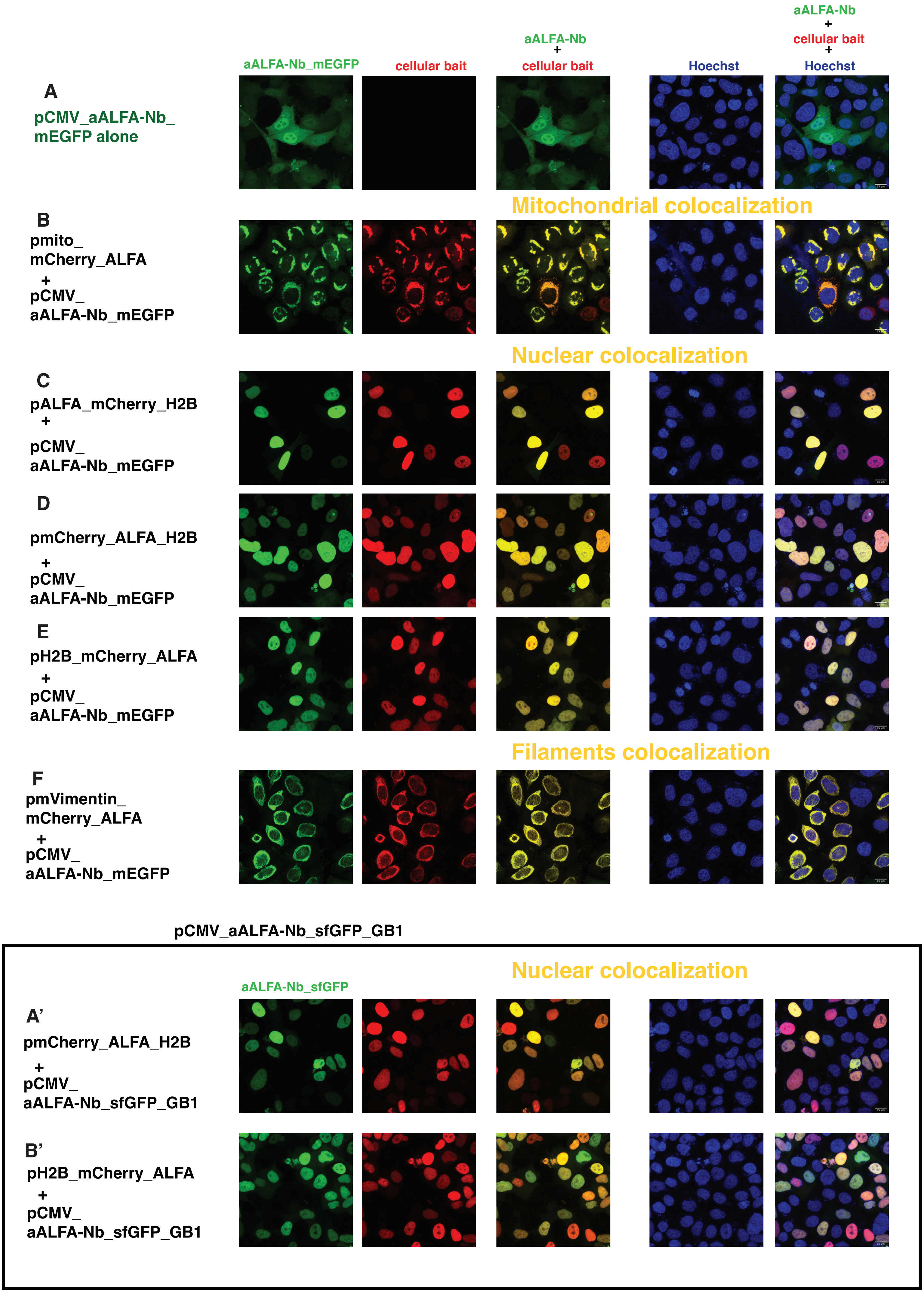


Vigano et al, Suppl. Figure8

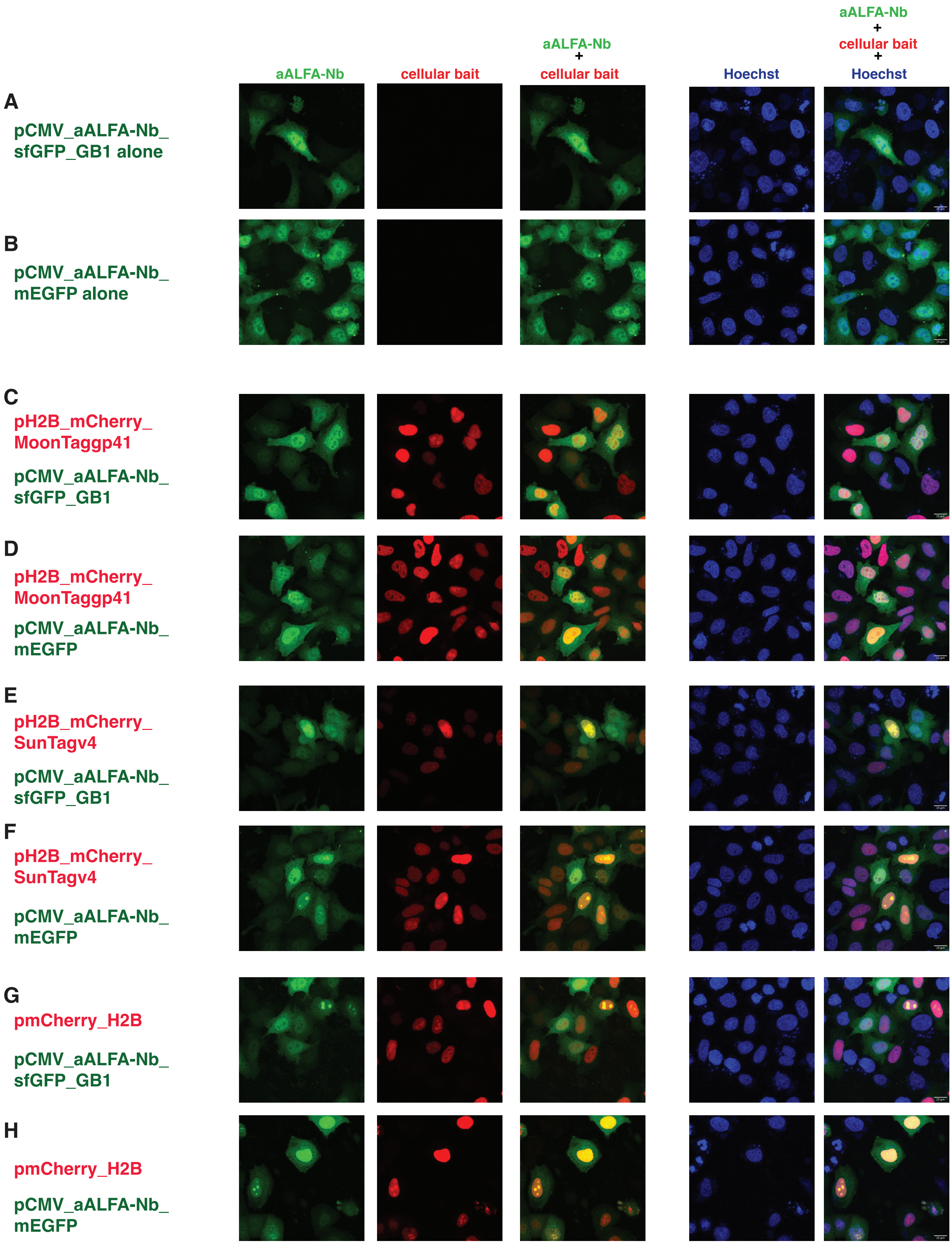


Vigano et al, Suppl. Figure9a

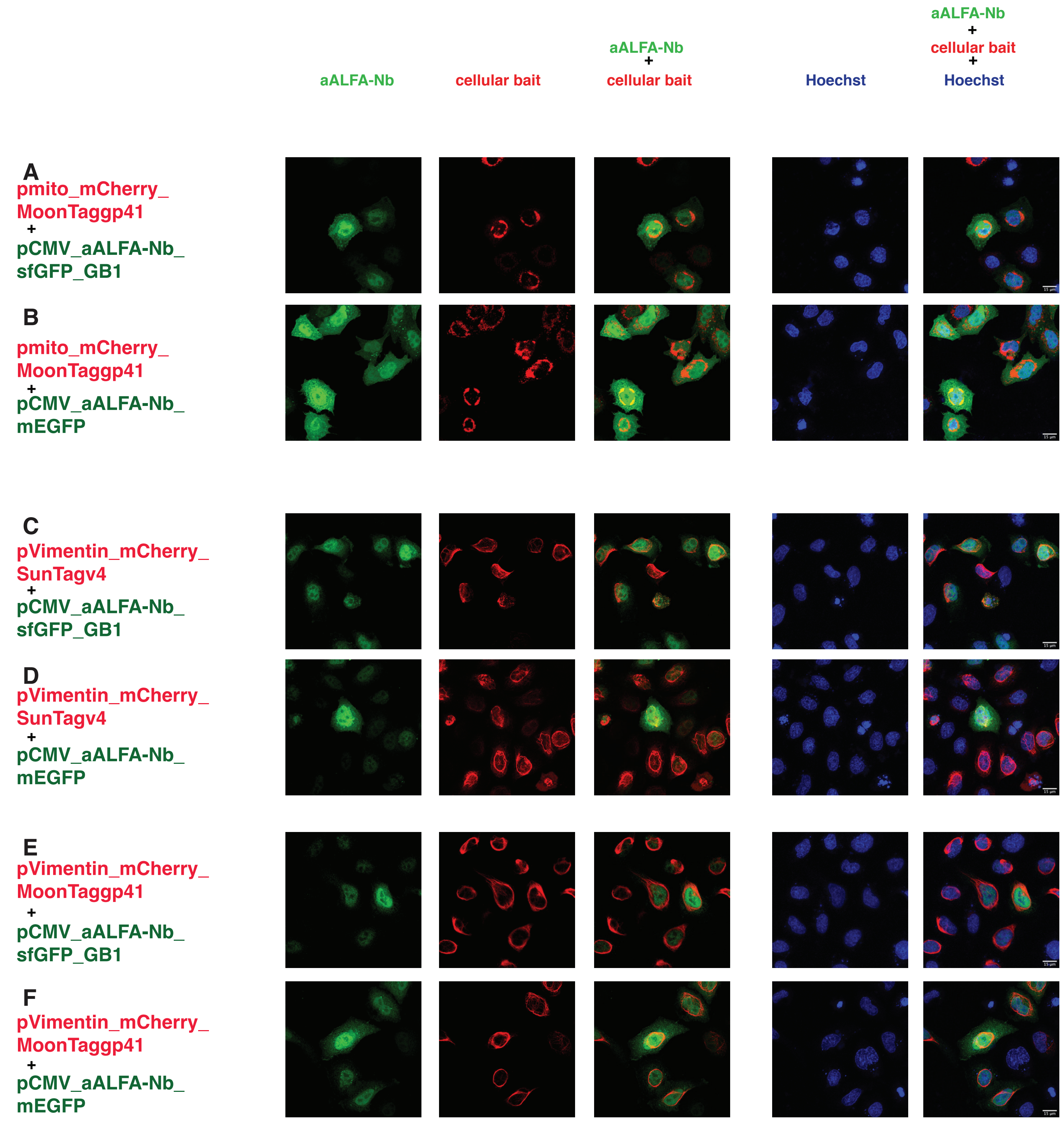


Vigano et al, Suppl. Figure9b

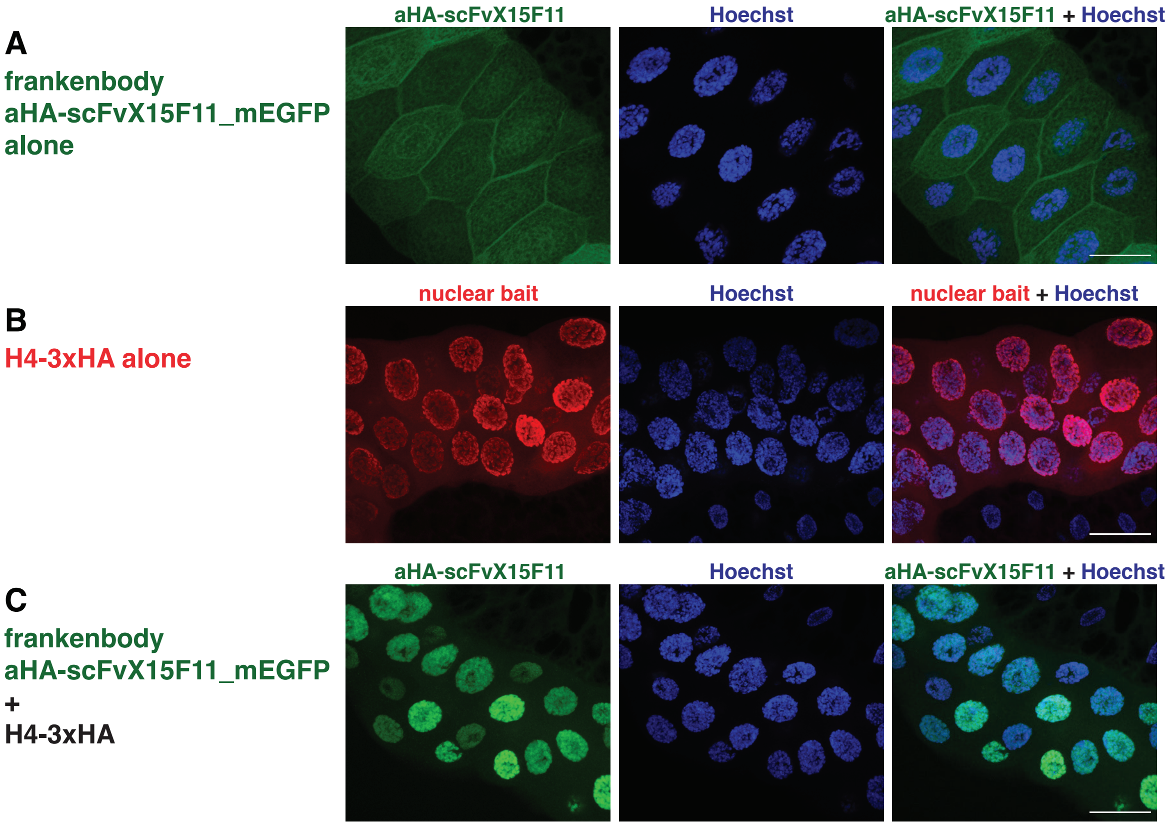


Vigano et al, Suppl. Figure10
